# A sequence context–unbiased approach to enable precise inference of mutational processes in cancer

**DOI:** 10.64898/2026.11.25.769148

**Authors:** Felicitas Bierth, Giulia Pizzolato, Christian Müller, Justinas Valiulis, Magdalena Seiffert, Filippo Beleggia, Alicja Paschowleska, Michael R. Schweiger, Marc R. Radke, Elisabeth Swisher, Matthias Fischer, Julie George, Roman Thomas, Robert Hänsel-Hertsch, Martin Peifer, Nima Abedpour, Maria Cartolano

**Affiliations:** Department of Translational Genomics, Medical Faculty, University of Cologne, Weyertal 115b, 50931 Cologne, Germany; Centre for Molecular Medicine, University of Cologne, Robert-Koch-Str. 21, 50931 Cologne, Germany; Department of Otorhinolaryngology, Head and Neck Surgery, Medical Faculty, University of Cologne, Kerpener Strasse 62, 50937 Cologne, Germany; Department I of Internal Medicine, Centre of Integrated Oncology Cologne-Bonn, University Hospital Cologne, Germany; Institute of Pathology, University of Cologne, Kerpener Str. 62, 50924 Cologne, Germany; Mildred Scheel School of Oncology Aachen Bonn Cologne Düsseldorf (MSSO ABCD), Faculty of Medicine and University Hospital Cologne, University of Cologne, Cologne, Germany; Institute for Translational Epigenetics, University Hospital Cologne, Faculty of Medicine, University of Cologne, Cologne, Germany; Cologne Centre for Genomics, CCG, Medical Faculty and Faculty of Mathematics and Natural Sciences, University of Cologne, Cologne, Germany; West German Genome Centre, WGGC, University of Cologne, Cologne, Germany; University of Washinton, Dept of Ob/Gyn, Division of Gynecologic Oncology, Seattle (WA), United States of America; Fred Hutchinson Cancer Centre, Clinical Research Division, Seattle (WA), United States of America; Department of Experimental Pediatric Oncology, University Children’s Hospital of Cologne, Medical Faculty, Cologne, Germany

## Abstract

Mutational signatures are recurring patterns of somatic mutations observed in cancer genomes, resulting from the activity of endogenous and exogenous mutagenic processes. Their investigation has significantly advanced our understanding of cancer etiology, informing prevention strategies and supporting clinical stratification. Although analytical approaches facilitate the identification of mutational signatures, their biological interpretation can be challenging, as some signatures may reflect methodological artifacts rather than distinct underlying mutational processes. Conventional mutational signatures inference frameworks generally assume that mutational opportunities are uniformly distributed across the genome and comparable across cancer types. However, mutable DNA motifs are not evenly distributed throughout the genome and their accessibility to DNA repair machinery is influenced by local chromatin states. To address these limitations, we developed *SEQUENCE* (*SEQ*uence context *U*biased *InferENCE* of mutational signatur*Es*), a method that accounts for the heterogeneous nucleotide composition by generating weighted mutation catalogues, in which each mutation is adjusted for the local availability of its respective sequence context. By using *SEQUENCE*-generated mutation matrices as input for *SigProfilerExtractor*, a conventional framework for mutational signature extraction, we demonstrate that our approach mitigates biases arising from cell-type specific and locally varying mutational rates across the genome. Furthermore, the latent local information encoded in the weights enables the disentanglement of co-occurring mutational processes targeting the same sequence context and refines the mutational spectra of previously described mutational signatures. Mutational signatures characterised by relatively flat mutational spectra are notoriously difficult to extract. Here, using clinically relevant mutational signatures, we demonstrate that *SEQUENCE* can improve patient stratification according to their response to adjuvant chemotherapy. Together, these findings highlight sequence context unbiased mutational signature inference as a promising new framework for revisiting current reference signatures, providing deeper insight into their underlying origins and facilitating their translation into cancer prevention and clinical application.

## Introduction

Cancer develops through the progressive accumulation of somatic genomic alterations generated by diverse endogenous and exogenous mutational processes^1^. Environmental exposures and lifestyle factors modulate the activity of many of these processes, shaping the mutational landscape of individual tumours and ultimately influencing cancer risk. Because each mutational process leaves characteristic patterns, or mutational signatures, in the genome, deciphering these genomic scars has become fundamental for understanding cancer etiology, identifying environmental carcinogens, and guiding therapeutic strategies^2,3,4,5^. Advances in high-throughput whole-genome sequencing over the past decades have enabled the systematic characterization of mutational signatures across human cancers, culminating in comprehensive reference catalogues of mutational processes^6,7^. A fundamental assumption underlying conventional mutational signature inference is that mutations observed across different genomic regions can be directly aggregated, implicitly treating the probability of observing a mutable DNA motif uniform across the genome. However, this assumption is not biologically realistic. Local mutation rates are influenced by multiple layers of genome organization. Repressive chromatin states, late replication timing, and low transcriptional activity correlates with high local mutation rate at the megabase scale. Whereas active transcription factor binding sites and transcription-coupled nucleotide excision repair anti-correlate with regional variation in mutation rate at the kilobase scale^8,9^. Disregarding the heterogeneity of these features across cancers with distinct (epi)genomic states, inflates the effect of tumour specific local mutation rates. Inference of mutational signatures independently for each tumour cohort minimises this bias^7^ but it does not address the regional heterogeneity along the genome. Recently, frameworks such as TensorSignatures^10^ or MuTopia^11^, albeit based on different assumptions, jointly modelled mutational spectra together with epigenomic features to simultaneously infer mutational signatures and their variable activity along the genome. Collectively, these studies emphasize the tight relation between local (epi)genomic states and the readout of the mutational processes. Nevertheless, these tools rely on non-cell of origin matched epigenomic data, thereby their applicability and their generalization across cohorts for *post hoc* analysis is limited.

In this study, we emphasise that the non-uniform nucleotide composition of the genome and the resulting heterogeneity in local sequence context availability represents an important consideration in mutational signatures studies and it is intrinsically coupled to higher-order genome organization. An illustrative example is provided by ultraviolet light-induced mutagenesis in skin melanoma, where the apparent distinction between the mutational spectra of signatures SBS7a and SBS7b, whose mutational spectra can be observed in Cosmic^12^, is not the result of two distinct mutational processes but rather a readout of the heterogeneous sequence context availability between early and late replicating regions, where these two signatures are known to be differentially active^13,14,15^.

To disentangle the intrinsic mutation spectrum of a damaging process from regional variation in genome sequence, *SEQUENCE*, or hereafter called also *SEQ*, creates weighted mutational catalogues, where each mutation implicitly carries the information of the local availability of the sequence context in which it is observed.

We applied *SEQUENCE* to 2638 whole-genome sequenced tumours from the ICGC/TCGA Pan-Cancer Analysis of Whole Genomes (PCAWG)^16^ project together with in-house whole-genome sequencing cohorts of small cell lung cancer (SCLC)^17^ and neuroblastoma (NB)^18^. Weighted mutation catalogues, constructed using the conventional 96-mutation classification, were analysed using *SigProfilerExtractor*, thus enabling direct comparison of the *SEQ*-based mutational signatures with well-characterised signatures inferred using conventional approaches. Using experimental cell lines exposed to cisplatin^19^, we demonstrate that *SEQ* mitigates the influence of cell-type specific difference during mutational signature inference. Similarly, in melanoma, *SEQ* reconstruct an intrinsic UV-mutational signature that is independent of regional mutational activity. Furthermore, it enables a clear distinction between the mutational process operating in cutaneous and acral melanoma, two subtypes with distinct etiological origin. In small cell lung cancer, *SEQ* disentangles co-occurring mutational processes associated with exposure to tobacco carcinogens, revealing additional complexity in smoking-induced DNA damage. Furthermore, *SEQUENCE*-derived signatures improve the identification of tumours with homologous recombination deficiency (HRD), enhancing patient stratification for adjuvant chemotherapy. Finally, analysis of memory B-cells and germinal centre-derived lymphomas identifies a mutational signature consistent with DNA polymerase η activity together with a prominent excess of C>T mutations at NCG trinucleotides, suggesting a contribution from enzymatically mediated deamination of 5-methylcytosine.

*SEQ* is implemented in Python and its source code will be provided alongside this manuscript. It is designed to enable a flexible choice of the genomic windows and the sequence context classification. Importantly, it requires only an opportunity genome (e.g. human, mouse) and the corresponding somatic mutation data to generate weighted matrices of mutation catalogues. These weighted catalogues can be then readily used with any matrix factorization framework for mutational signature extraction. By accounting for local availability of mutable sequence contexts, *SEQ* aims to provide a more accurate representation of the mutational processes shaping the cancer genomes.

## Results

### Local triplet-density correction with *SEQ* reduces cell-type-associated variation in inferred signatures of cisplatin activity

Somatic mutation rates vary considerably along the genome and are influenced by multiple layers of genomic organization^20^. The nucleotide composition of the genome itself imposes heterogeneity of the local sequence context mutability (Suppl. Fig.1a, 1b). Because of this difference in trinucleotide frequencies between genomic regions (Fig. 1a, 1b), a mutation at a locally rare trinucleotide reflects a higher per-site rate than the same mutation where that trinucleotide is common (Fig. 1c). Thus, the common assumption that the mutational opportunity of a DNA sequence context is equal along the genome, biases mutational signature inference. We reason that local sequence context is intrinsically coupled to higher order of genome organisation^13,14^. Consequently, mutational signatures inferred without accounting for local trinucleotide opportunity in different tissues may be affected by cell-type-associated mutational spectra^21^. Indeed, one mutational process acting in two cell types yields distinct raw spectra (Fig. 1d), whereas weighting each mutation by the inverse of the local frequency of its trinucleotide recovers the same intrinsic per-site signature in both (Fig. 1e). To establish whether *SEQ* correction reduces this bias, we re-analysed cisplatin exposure data generated in two cell lines of distinct developmental origin: Hep2G, a liver cancer cell line, and MCF10A, a non-tumourigenic breast epithelial cell line^19^. Signature extraction following *SEQ* correction using four Hep2G and six MCF10A replicates identified two signatures as the optimal solution (Fig. 2a, Suppl. Fig. 2a, 2c). The first, *SEQ*-Ciplatin-SBS96A, contributed comparably across the cell lines (Fig. 2a, 2c) and displayed the canonical cisplatin-associated damage, namely C>T, T>A and C>A. The second, *SEQ*-Ciplatin-SBS96B, was predominantly associated with MCF10A replicates 4 and 5 and was consistent with a possible technical artefact introduced during sample preparation (Suppl. Fig. 2e). On the contrary, signature extraction from uncorrected mutation counts identified four signatures as the optimal solution (Fig. 2b and Suppl. Fig. 2b, 2d). One signatures, *Counts*-Cisplatin-SBS96C, was consistent with the putative technical artefact also described above (Fig. 2b and Suppl. Fig. 2e). The remaining three signatures showed similar mutation profiles, all consistent with canonical cisplatin damage^19^, but differed in their activities between the two cell types (Fig. 2b, 2d), suggesting that signature inference from uncorrected counts separated the cisplatin-associated profile into cell-type-associated biases of mutational composition.

**Fig. 1:**
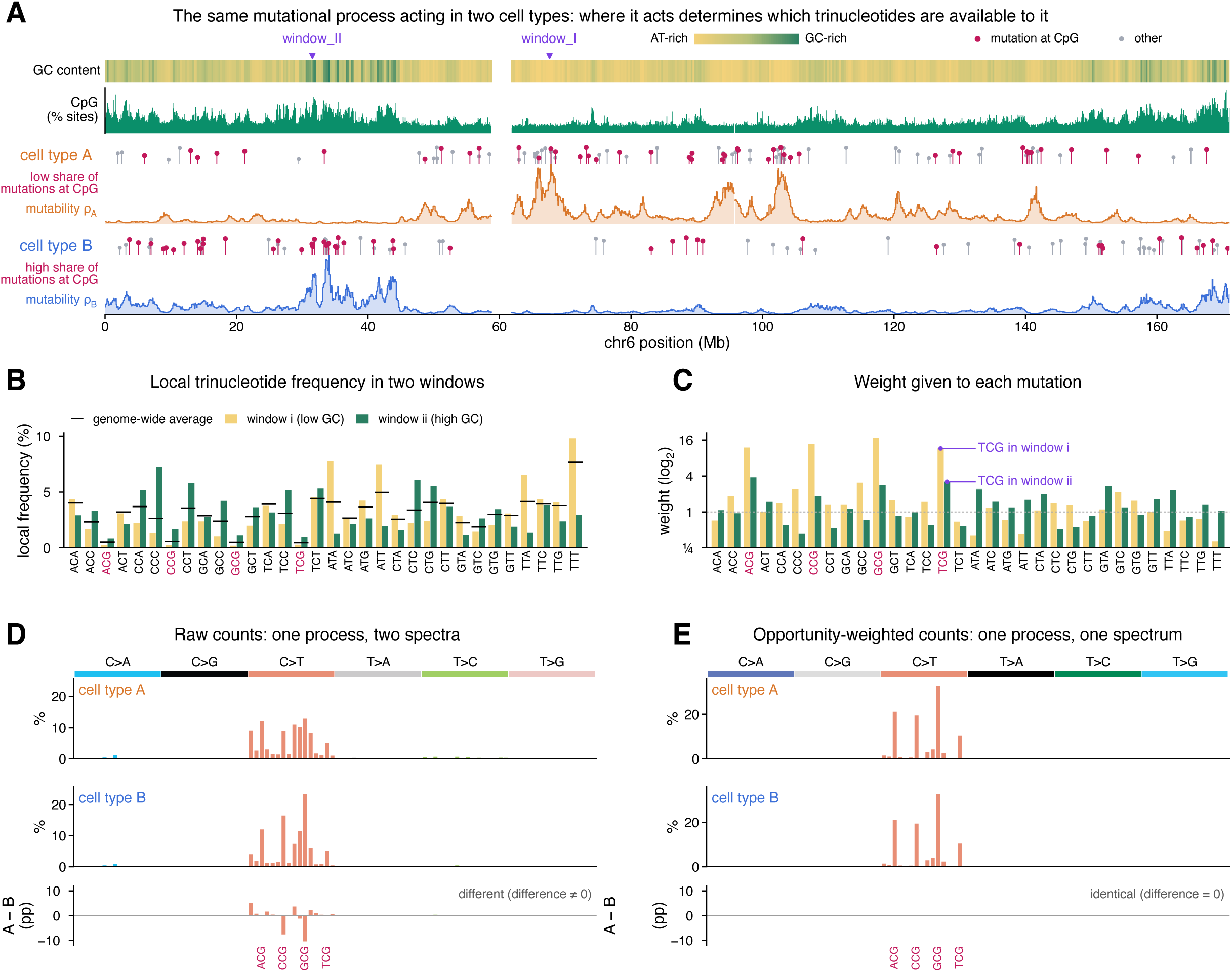
**Opportunity-weighted mutation counts recover cell-type-invariant mutational signatures**: A) One mutational process acting in two cell types along human chromosome 6. The top tracks show local GC content and CpG frequency; gaps mark the centromere and unassembled regions. Cell type A preferentially mutates AT-rich DNA and cell type B GC-rich DNA (regional mutability ρ_A and ρ_B, illustrative). Lollipops show a random sample of mutations in each cell type, with mutations at CpG in magenta; the share of mutations at CpG is lower in A than in B. Arrows (indicated as window_I and window_II) mark two exemplary genomic windows. B) Local frequency of each trinucleotide in a low-GC (window i) and a high-GC (window ii) window, with the genome-wide average shown as black ticks. C) The same TCG mutation weighs more in the low-GC window_I, where TCG is rare, than in the high-GC window_II. D) Raw 96-channel catalogues of the same process in cell types A and B, which differ, and their difference in percentage points. E) Opportunity-weighted catalogues of the same samples, which are identical: both equal the intrinsic per-site signature of the process, and their difference is zero.

**Fig. 2:**
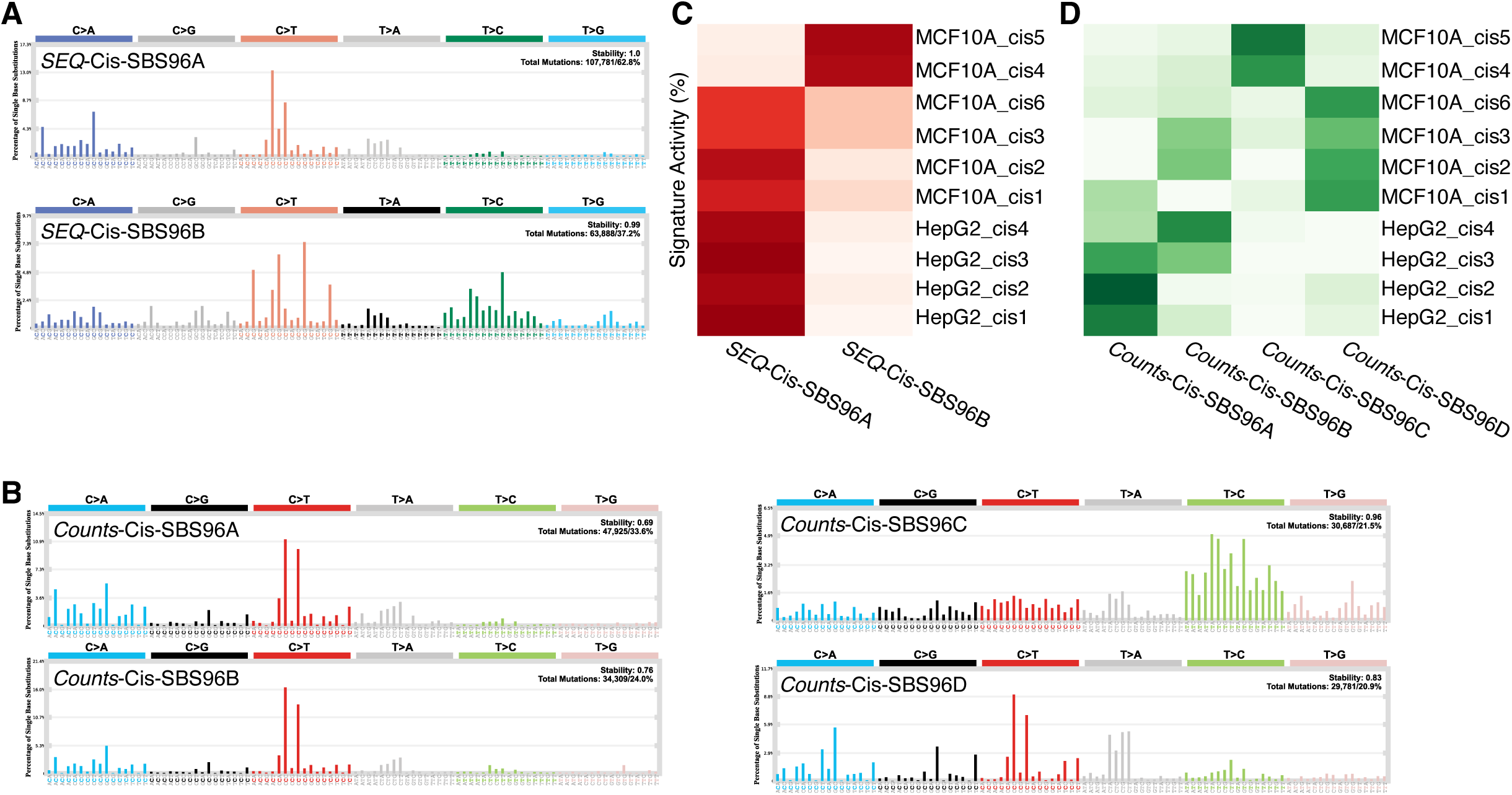
***SEQ* mitigates the influence of cell type-specific local mutation rates:** A) *SEQ*-derived mutational signatures of cisplatin exposure (*SEQ*-Cis). B) *Counts*-derived mutational signature of cisplatin exposure (*Counts*-Cis). C) *SEQ*-Cis and D) *Counts-*Cis exposures in MCF10A (replicate 1 to 5) and HepG2 (replicates 1 to 4) cell lines.

Together, these results support the hypothesis that correcting for local trinucleotide density with *SEQ* can reduce the bias of cell-type-associated differences in mutation distribution along the genome and thereby in mutational signature inference. In this experimental setting, in which both cell lines were exposed to the same chemical agent, *SEQ* correction recovered a shared cisplatin-associated profile rather than multiple signatures reflecting cell-type specific regional variation in mutational opportunity.

### Mutational signatures caused by exposure to UV light

Solar ultraviolet radiation (UVR) is a potent genotoxic agent and the principal environmental cause of skin cancers. UV exposure induces covalent linkages between adjacent pyrimidines (cytosine or thymine), generating predominantly cyclobutane pyrimidine dimers (CPDs) and pyrimidine-6,4-pyrimidone photoproducts (6,4PPs)^22^. Cytosines within these unstable dimers are prone to spontaneous deamination to uracil, which, if not promptly repaired by the nucleotide excision repair machinery (NER), is ultimately replicated as a thymine^6^.

*SEQ-*based mutational signature inference in the skin melanoma cohort extracted *SEQ*-SkinMelanoma-SBS96A, a signature dominated by C>T substitutions at dipyrimidine contexts (notably <u>T[C</u>>T]N and <u>C[C</u>>T]N). This signature is predominantly active in the skin cutaneous melanoma subtype, consistent with its attribution to UVR (Fig. 3a, 3b). Differential trinucleotide availability between early- and late-replicating regions is a recognised source of bias for the inference of the UVR mutational signature and it is known to contribute to the mutational profile of SBS7a and SBS7b (Suppl. Fig. 3a, 3b)^10,13,14^. In line with our hypothesis, *SEQ*-based correction of local triplet availability removed this bias and consequently returned a consensus mutational signature associated to UVR damage.

**Fig. 3:**
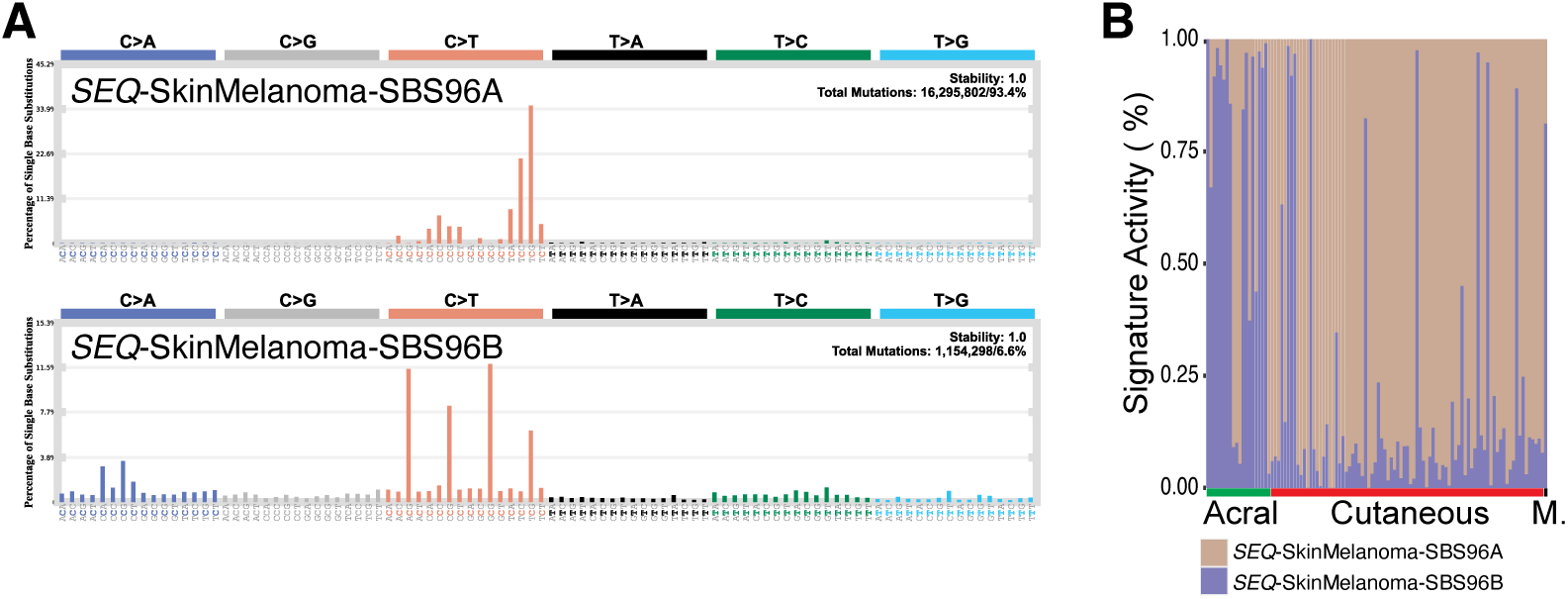
***SEQ*-derived mutational signatures extraction in the skin melanoma cohort.** A) *SEQ*-derived mutational signatures profiles (*SEQ*-SkinMelanoma). B) Stacked bar graph of relative mutational signature activity in the skin melanoma cohort. The three different subtypes are indicated below: acral, cutaneous and mucosal melanoma (M.).

Furthermore, we observed a conspicuous peak at TCG sites in the *SEQ*-SkinMelanoma-SBS96A profile (Fig. 3a). Facilitated CPD formation has been observed at 5-methyl cytosine (5-meC) of CpG sites^23^ and, UV light-induced mutation rate of TCG triplets is reduced in active promoters^24^. Thus, *SEQ-*based mutational signature extraction infers a mutational spectrum of UVR damage that is affected neither by the availability of the sequence context nor by its visibility to the DNA repair machinery.

A second mutational signature inferred in the Skin Melanoma cohort, *SEQ*-SkinMelanoma-SBS96B, is characterized predominantly by C>T substitutions at CpG sites and displays preferential activity in the acral and the mucosal subtype. Its distribution across subtypes and its limited association with the cutaneous subtype argues against UVR as its mutagenic source (Fig. 3a, 3b). Albeit *SEQ*-SkinMelanoma-SBS96B mutation pattern recalls the spectrum of mutations resulting from the deamination of the 5-meC, its activity is not associated with patient age (robust linear regression, p-value =0.22), therefore it is unlikely the result of an age-dependent accumulation of somatic mutations^25^. The preferential activity of *SEQ*-SkinMelanoma-SBS96B in tumours carrying the *BRAF* V600E mutant allele and its conspicuous pattern is rather suggestive of AID-mediated 5-meC deamination as the underlying mutational process (Suppl. Fig. 3c)^26^. However, these observations need further validation.

Taken together, *SEQ*-based inference of mutational signatures allows the extraction of a spectrum reflecting the UVR mutation process unbiased from (epi)genomic layers of the genome and it allows for a clear separation between tumor subtypes of distinct etiologies.

### The mutational signatures caused by exposure to cigarette smoke

Polycyclic aromatic hydrocarbons (PAHs) are within the most abundant carcinogenic compounds found in cigarette smoke (CS)^27^. They form bulky DNA adducts at guanine residues, which if not repaired by transcription coupled nucleotide excision repair (TC-NER), can lead to C>A/G>T transversion upon DNA replication^27^. Furthermore, benzo[a]pyrene -particularly its active metabolite, benzo[a]pyrene diol epoxide (BPDE) -binds with high affinity to methylated CpG regions, forming DNA adducts at guanine residues^28^. In line with these observations, C>A/G>T transversions caused by polycyclic aromatic hydrocarbons occur more frequently at methylated CpG dinucleotides *in vitro* and have been frequently observed in small cell lung cancer^29,30,31^. In addition, PAHs can induce oxidative stress via the generation of reactive oxygen species (ROS)^32^. ROS can damage the DNA by oxidazing guanine to form 8-oxo-7,8-dihydroguanine (8-oxoG). If this lesion is not repaired by the base excision repair (BER) pathway, it can likewise result in to C>A/G>T transversions during DNA replication. The signature extraction of these processes can be therefore hindered by the fact that both mutational processes can target the same nucleotide in the sample.

The property of *SEQ* is to assign a weight to each triplet inversely proportional to its local availability. An evident consequence of this effect, is that NCG triplets, otherwise sparse in the genome, can acquire larger relative values to influence signature extraction (Suppl. Fig. 1). In line with this hypothesis, *SEQ*-based mutational signature inference in SCLC, Lung Adenoca, Lung-SCC, Head-Neck (the HNSC-US cohort, only smokers) and the Liver-HCC cohort resulted in to CS-associated mutational signatures that could be separated in to two distinct branches upon hierarchical clustering of pairwise cosine similarities (Fig. 4a, Suppl. Fig. 4). Conversely, tree topology of CS-induced *Counts* signatures did not lead to similar interpretations with multiple signatures inferred from the same tumour type clustering within the same clade (Suppl. Fig. 5a).

**Fig. 4:**
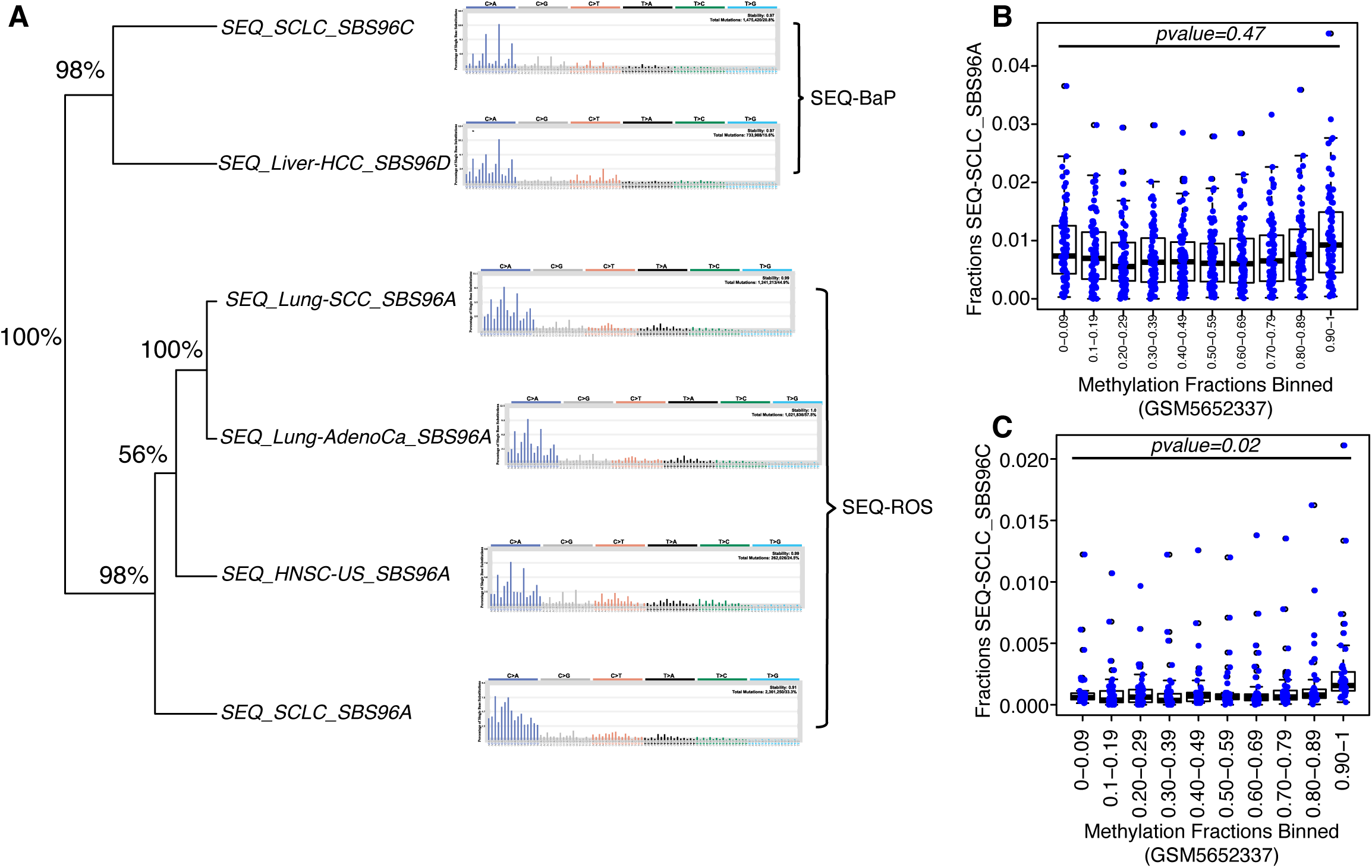
***SEQ*-derived mutational signatures of cigarette smoke exposure.** A) Hierarchical clustering of *SEQ*-derived mutational signatures resulting from the exposure to cigarette smoke. Branch support is obtained from multinomial resampling of the original mutational catalogues and it is shown as percentage on the left of the tree. The signatures are separated in two clades, *SEQ*-BaP and *SEQ*-ROS, based on their putative etiology. B) Fractions of mutations assigned to *SEQ*-SCLC_SBS96A and to C) *SEQ*-SCLC_SBS96C at methylated CpG sites measured in a lung bronchial tissue cell lines from healthy donors (GSM5652337). Methylation fractions are separated in to bins. P-values obtained from student’s t-test comparing fractions in the first (0-0.09) and the last bin (0.90-1) are indicated on the top of the boxplots.

We postulate that these two branches may represent the main mechanisms of DNA damage associated with exposure to CS in these cancers: one likely induced by the pro-oxidant components present in CS (Fig. 4a, clade *SEQ*_Tobacco_ROS), the other induced by the formation of DNA adducts at guanine bases within methylated CpG regions (Fig. 4a, clade *SEQ*_Tobacco_BaP). The *SEQ*_Tobacco_ROS branch is populated by mutational signatures inferred in lung tumours, likely recapitulating the activity of the canonical tobacco signature SBS4 in Cosmic. The *SEQ*_Tobacco_BaP branch is populated by a mutational signature found in Liver-HCC (*SEQ*_Liver_HCC_SBS96D) and one found in SCLC (*SEQ*_SCLC_SBS96C) (Fig. 4a) and it is characterized by C>A substitutions at NCpG trinucleotides. We found that mutations attributed to *SEQ*_SCLC_SBS96C were enriched at methylated sites identified in a lung bronchial tissue from healthy donors^33^ (Fig 4b , 4c, Suppl.Fig. 5b-5e) suggesting that these mutations can be the result of oxidation of the guanine residue neighbouring 5-meC.

**Fig. 5:**
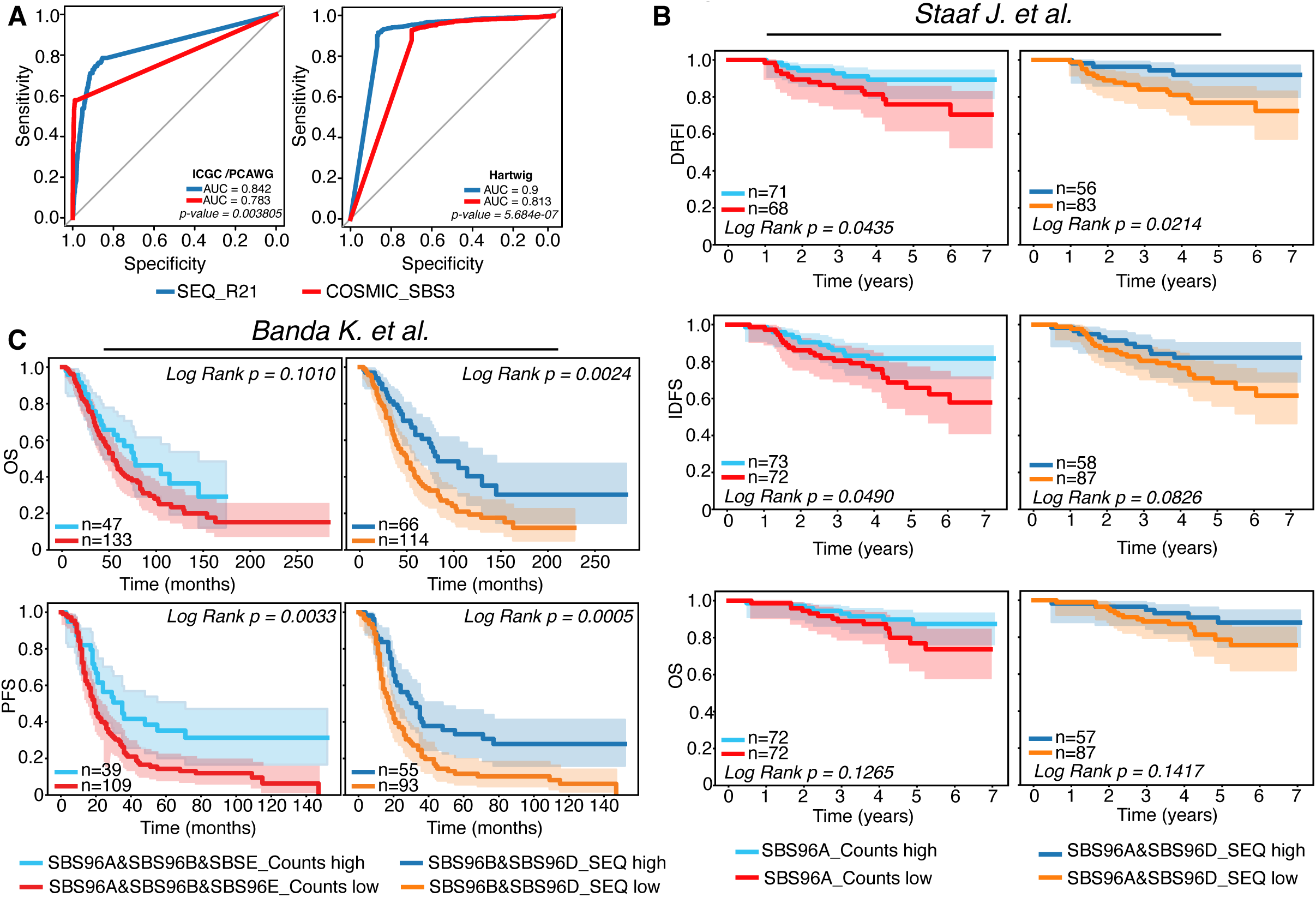
***SEQ*-derived mutational signatures improve the prediction of HRD tumours:** A) ROC curve and AUC reflecting the accuracy of *SEQ*_R21 (blue) and Cosmic_SBS3 (red) in predicting HRD across the ICGC/PCAWG (left) and the Hartwig (right) cohort. B) Kaplan Meier curves and log rank test reflecting the ability of HRD-associated mutational signatures to stratify patients by DRFI, IDFS and OS in the triple negative breast cancer cohort published in *Staaf et al.*, C) Kaplan Meier curves and log rank test reflecting the ability of HRD-associated mutational signatures to stratify patients by OS and PFS in the tubo-ovarian carcinoma Washington cohort published in *Banda et al.* DRFI: Disease Relapse Free Interval, IDFS: Invasive Disease Free Survival, OS: Overall Survival, PFS: Progression Free Survival. Red and light blue indicate patients stratified according to the activity of the HRD signatures estimated with raw counts as input. Orange and dark blue indicate patients stratified according to the activity of the HRD signatures estimated with *SEQ*-weighted mutational catalogues as input. High and Low indicate above/equal and below median signature activity, respectively. n= number of patients belonging to the above and below the median groups, respectively. SBS96A_Counts indicate patients stratified by the activity of SBS96A. SBS96A&SBS96D_*SEQ* indicate patients stratified by the activity of SBS96A and SBS96D. SBS96A&SBS96B&SBS96E_*Counts* indicate patients stratified by the activity of SBS96A and SBS96B and SBS96E. SBS96B&SBS96D_*SEQ* indicate patients stratified by the activity of SBS96B and SBS96D.

Thus, incorporating mutation weighting with *SEQ* prior mutational signature inference enables the disentanglement of concurrent mutational processes acting on the same trinucleotide contexts, thereby providing deeper insights in to the mechanisms by which cigarette smoke induces DNA damage.

### The mutational signatures of Homologous-Based Repair Deficiency

Double DNA strands breaks are repaired by the Homologous Recombination (HR) repair machinery during the S and the G2 phase. Pathogenic mutations in the genes regulating HR have been mostly observed in breast, ovarian, pancreatic, and stomach cancers and make these cancers susceptible to DNA damaging chemotherapies (e.g., platinum) and to poly (ADP-ribose)^34^. Therefore, the ability to correctly distinguish HR-deficient (HRD) from HR-proficient (HRP) tumors is clinically relevant^5,35^. Cosmic SBS3 is a mutational signature often associated to HRD tumours and is thought to be caused by translesion synthesis over double strand breaks^6^. The activity of the SBS3 is alone insufficient to distinguish HRD from HRP tumours^36^, but it represents an important feature included in HRD classifiers^5^. The non-distinctive mutational profile of SBS3 is likely the cause of its aspecific assignment in HRP tumours^37^. Therefore, we sought to evaluate whether *SEQ* correction of mutation catalogues could improve mutational signature inference and activity assignments in HRD tumours. To test our hypothesis, we initially assembled a consensus HRD signature, *SEQ*_R21, from the mean mutation spectrum of a cluster consisting of signatures associated to HRD (see Fig.2 and methods). Next, we tested whether *SEQ*_R21 activities could predict HRD tumours and, if so, with higher discriminative performance than SBS3. In line with our hypothesis, *SEQ*_R21’s activities showed an improved predictive performance than SBS3’s both in the ICGC/PCAWG and in the Hartwig cohorts (Fig. 5a).

Furthermore, we evaluated whether *SEQ*-derived signatures associated to HRD could support patient stratification by treatment outcome (Fig. 5b, 5c, Suppl. Fig. 6 and Suppl. Fig. 7). To this end, we analysed the Sweden Cancerome Analysis Network-Breast (SCAN-B) cohort of triple negative breast cancers^5^ and the University of Washington cohort of tubo-ovarian carcinomas^35^. Mutational signatures were extracted both from *SEQ*-derived and conventional counts mutational catalogues. Association of mutational signatures with HRD was tested by means of a logistic regression (see methods, Fig.5 and Suppl. Fig. 6 and Suppl. Fig. 7).

In breast we observed that patients exhibiting concomitantly high activities of SBS96A_*SEQ* and SBS96D_*SEQ* showed significantly improved distant relapse-free interval DRFI (DRFI, log-rank P=0.0214, Fig. 5b top right panel, Suppl. Table 4). This association was more statistically significant than that observed for SBS96A_*Counts* alone (log-rank P=0.0435, Fig. 5b top left panel, Suppl. Table 5). On the contrary, concomitant high activities of SBS96A_*SEQ* and SBS96D_*SEQ* did not significantly stratify patients according to invasive disease-free survival (IDFS, log-rank P=0.0826, Fig. 5b, middle right panel), whereas high SBS96A_*Counts* activity allowed only a marginal separation between the two groups (log-rank P=0.0490, Fig. 5b middle left panel). Finally, stratification based on the activity of the HRD signatures suggested a trend towards improved overall survival (OS), however, neither association reached statistical significance (log-rank P=0.1265 and log-rank P=0.1417, Fig. 5b bottom left and right panels, respectively).

In ovarian carcinoma, we extracted two *SEQ*-based signatures associated to HRD (Suppl. Table 6, Fig. 5c) and patients with the combined high activities of both showed improved OS and progression free survival (PFS) upon standard of care platinum chemotherapy (Fig. 5c, right panels, log-rank P=0.0024 and log-rank P=0.0005, respectively). Interestingly, high combined activities of HRD-associated signatures extracted from the conventional counts mutational catalogues could not separate patients as effectively (Suppl. Table 7, Fig. 5c left panels, log-rank 0.1010 and log-rank 0.0033).

These findings indicate that correcting for mutational opportunity enhances the inference of mutational signatures with flat, non-distinctive mutation spectra. This is particularly relevant when multiple mutational processes co-occur, as the stronger process may mask the activity of the weaker one, thus hindering its correct identification.

Importantly, we acknowledge that the limits posed by the lack of specificity of the HRD-associated signatures still remain with *SEQ* and that the activity of *SEQ*-derived HRD-associated mutational signatures cannot be used alone to accurately distinguish HRD from HRP tumours (Fig. 5a). However, our observations are suggestive of an effective improvement when correcting for mutational opportunity, thereby flagging the necessity of taking this correction in to account in the development of predictive models of HRD that utilise mutational signatures.

### The mutational signature of activation-induced cytosine deaminase in B-cells

Maturing B-cells in the germinal centre (GC) undergo genetic and functional changes that ultimately lead to the production of B-cells presenting high affinity antibodies. These processes are primarily mediated by the activation-induced deaminase (AID) enzyme that initiates somatic hypermutation (SHM), in the dark zone, and class switch recombination (CSR), in the light zone of the germinal centre, of the immunoglobulin genes^38^. To further sustain SHM and guarantee B-cell diversification, an error prone mismatch repair recognises and repair the damage induced by AID while introducing additional mutations at neighbouring nucleotides^39^. Beyond its activity on the immunoglobulin genes, AID catalyses the deamination of the 5-meC in maturing B-cells undergoing profound genome-wide methylation changes^40^.

In the dark zone, AID preferentially deaminates cytosines at WR<u>C</u>Y motifs of single-stranded DNA during transcription (W=A/T; R=A/G; Y=C/T) and it catalyses the deamination of cytosine to uracil. Failing to recognise the uracil prior DNA replication, results in to C>T transitions^41,42,43^. "Off target" AID activity has been shown to mutate genes and their associated regulatory elements influencing lymphomagenesis^44^. Accordingly, the mutational spectra of AID-mediated SHM at WR<u>C</u>Y DNA motifs is exquisitely reconstructed with the *SEQ*-based signature extraction in DLBCL active super enhancers, whereas interestingly, the *Counts*-based signature extraction reconstructs a profile with preference for WGCY DNA motifs of AID binding (Fig. 6a, b).

**Fig. 6:**
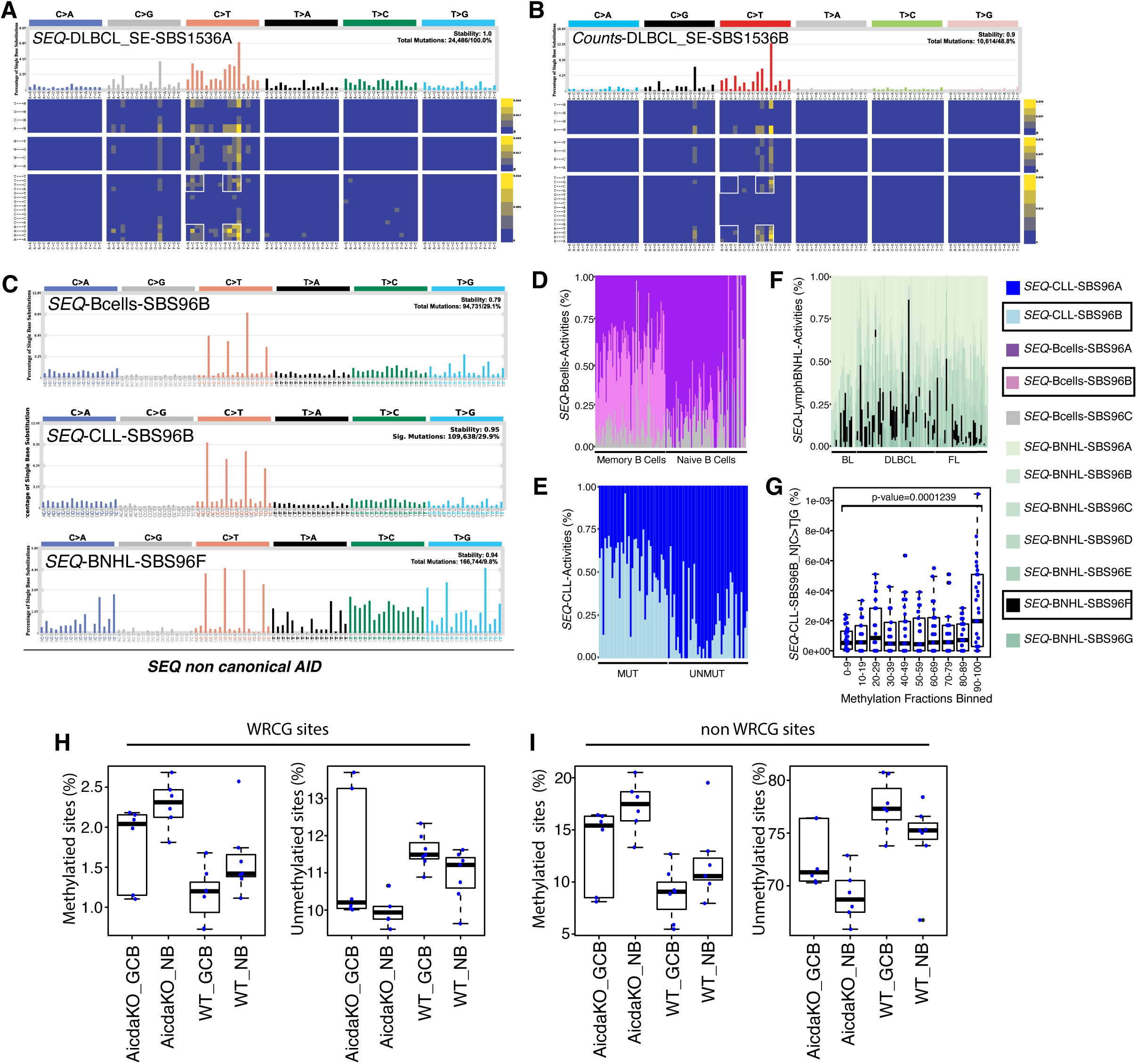
Mutational signatures of canonical and non-canonical AID in B-cells. SBS1536 mutational signature inference of DLBCL’s super enhancers using A) *SEQ*-weighted and B) raw *Counts* as input. White framed square highlight mutation probabilities at the WRCG context. C) *SEQ*-based mutational signatures of non-canonical AID independently inferred in a cohort of healthy memory and naive B-cells, Lymph-CLL and Lymph-BNHL tumours. Estimated exposure of the non-canonical AID signature in B-cells D), Lymph-CLL E), and Lymph-BNHL tumours F). The mutational signatures reflecting the non-canonical AID activity are indicated in a box within the label colour legend. G) Boxplots showing the association of the N[C>T]G mutations assigned to *SEQ*-CLL_SBS96C to methylation allelic frequencies in the lymphoblastoid cell line GM12878. P-value of a student’s t-test between bins (0-9) and (90-100) is indicated on top. H) Percentage of methylated (left panel) and unmethylated sites (right panel) at WRCG DNA motifs in wild type mice naive B-cell (WT_NB), wild type germinal centre B-cells (WT_GCB), naive B-cells in *Aicda* knockout (AicdaKO_NB) and germinal centre B-cells in *Aicda* knockout (AicdaKO_GCB). I) Percentage of methylated (left panel) and unmethylated sites (right panel) at non-WRCG DNA motifs in wild type mice naive B-cell (WT_NB), wild type germinal centre B-cells (WT_GCB), naive B-cells in *Aicda* knockout (AicdaKO_NB) and germinal centre B-cells in *Aicda* knockout (AicdaKO_GCB). MUT indicates Lymph-CLL tumours originating from the germinal centre. Whereas, UNMUT indicates Lymph-CLL tumours originated before entering the germinal centre. FL is Follicular Lymphoma, BL is Burkitt Lymphoma and DLBCL is Diffuse Large B-cell Lymphoma.

The polymerase η, lacking exonuclease activity and recruited by the mismatch repair machinery at resected DNA strands, synthesise DNA by adding mismatches at A and T nucleotides^39^. Accordingly, the Cosmic SBS9 signature, otherwise called non-canonical AID signature, shows a mutational spectrum with prevalence of T>C and T>G mismatches in a NTpW context^45^. While we could reconstruct a profile similar to SBS9 using *Counts*-based matrices (Suppl. Fig. 8a), *SEQ*-based reconstruction infers a mutational signature of non-canonical AID which is characterised by both C>T transitions at CpG sites and T>C and T>G mutations in a TpW context. We postulate that this mutational spectrum likely reflects the mutagenic activity both of polymerase η and of the AID-mediated deamination of 5-meC in the germinal centre (Fig. 6c). Several lines of evidence support our interpretation: *SEQ* non-canonical AID mutational signature is extracted independently in three cohorts: the Lymph CLL and the Lymph BNHL B-cell lymphoma cohorts of the ICGC/PCAWG consortium and the healthy single cell colonies-derived B-cells (Fig. 6c)^45^, thereby arguing against a cohort-specific mutational signature. The activity of these signatures is confined to the germinal centre B-cells (Fig. 6d, 6e, 6f) where they operate both on the immunoglobulin genes and genome wide (Suppl Fig. 8c-8h). Furthermore, C>T mutation at CpG sites assigned to the non-canonical AID signatures are enriched at methylated sites of a lymphoblastoid cell line representing mature B lymphocytes (Fig. 6g, Suppl. Fig. 8b). AID-mediated deamination of the 5-meC does not specifically target WRCG DNA motifs (Fig. 6h, 6i)^40^, therefore it is likely that these mutations are left unrepaired during AID-mediated deamination (Suppl. Fig. 9). Thus, *SEQ*-based extraction of mutational signatures in lymphoma and B-cells aids the inference of a non-canonical AID mutational process reflecting the concerted activity of the error-prone polymerase η and of the of 5-meC deamination, possibly mediated by AID itself. Whether these processes are mechanistically connected or whether they co-occur, is not possible to infer here. Our findings however speak for an additional mutational process, undescribed so far, shaping B-cells in the germinal centre.

### Establishing a reference set of *SEQ*-based mutational signatures

*SEQ* supported the inference of 127 mutational signatures across 2796 samples and 32 cohorts. The *SEQ*-based mutational signatures reconstructed the original mutation catalogues with an accuracy comparable to that achieved with the conventional method (Suppl. Fig. 10b, 10c). By performing hierarchical clustering of mutational signatures pairwise cosine similarities, we established an initial reference set consisting of 21 clusters and 10 singletons (Fig. 7, Suppl. Fig. 10a and Suppl. Fig. 11a). Based on pattern similarity with the available Cosmic dataset and on the preponderance of signatures exposure in specific tumour cohorts^6^, we could confidently assign *SEQ*-based reference signatures to exogenous sources of damage such as aristolochic acid (*SEQ*_R01) or exposure to sunlight (*SEQ*_S04). Furthermore, as referred early in the manuscript, we distinguish two clusters of mutational signatures induced by cigarette smoke (*SEQ*_R03 and *SEQ*_R04). Endogenous sources of damage were assigned to *SEQ*_R05 (similar to SBS18, damage by reactive oxygen species), *SEQ*_R06 (similar to SBS10ba and SBS10b, loss of proofreading activity of pol epsilon), *SEQ*_R07 (similar to SBS12 and SBS13, APOBEC activity), *SEQ*_R15 (similar to SBS1, spontaneous cytosine deamination), *SEQ*_R20 (similar to SBS92, tobacco smoking and damaging mutations in *ERCC2*), SEQ_R21 (similar to SBS3, HRD), *SEQ*_S03 (similar to SBS2, APOBEC activity), and SEQ_S09 (similar to SBS9 with additional C>T alterations at NCG triplets, non-canonical AID). *SEQ*-based mutational signatures with unknown etiology in Cosmic are *SEQ*_R02 (similar to SBS17a and SBS17b), *SEQ*_S02 (similar to SBS12) and *SEQ*_S06 (similar to SBS11). We also report *SEQ*_R17, a signature observed in gynecological cancers of the ICGC/PCAWG cohort, i.e. Uterus-AdenoCa, Ovary-AdenoCa and Breast-AdenoCa. This signature is characterised by the prevalence of C>A, C>G and C>T substitutions at CpG sites and its activity is associated to HRD in breast and uterus adenocarcinoma samples (Suppl. Table 3). To our knowledge, a signature displaying a similar mutation spectrum has not been previously identified in the ICGC/PCAWG cohort. However, a signature with a similar mutation pattern has been reported in one bladder and one lung cancer sample of the Genomics England (GEL) cohort^7^.

**Fig. 7:**
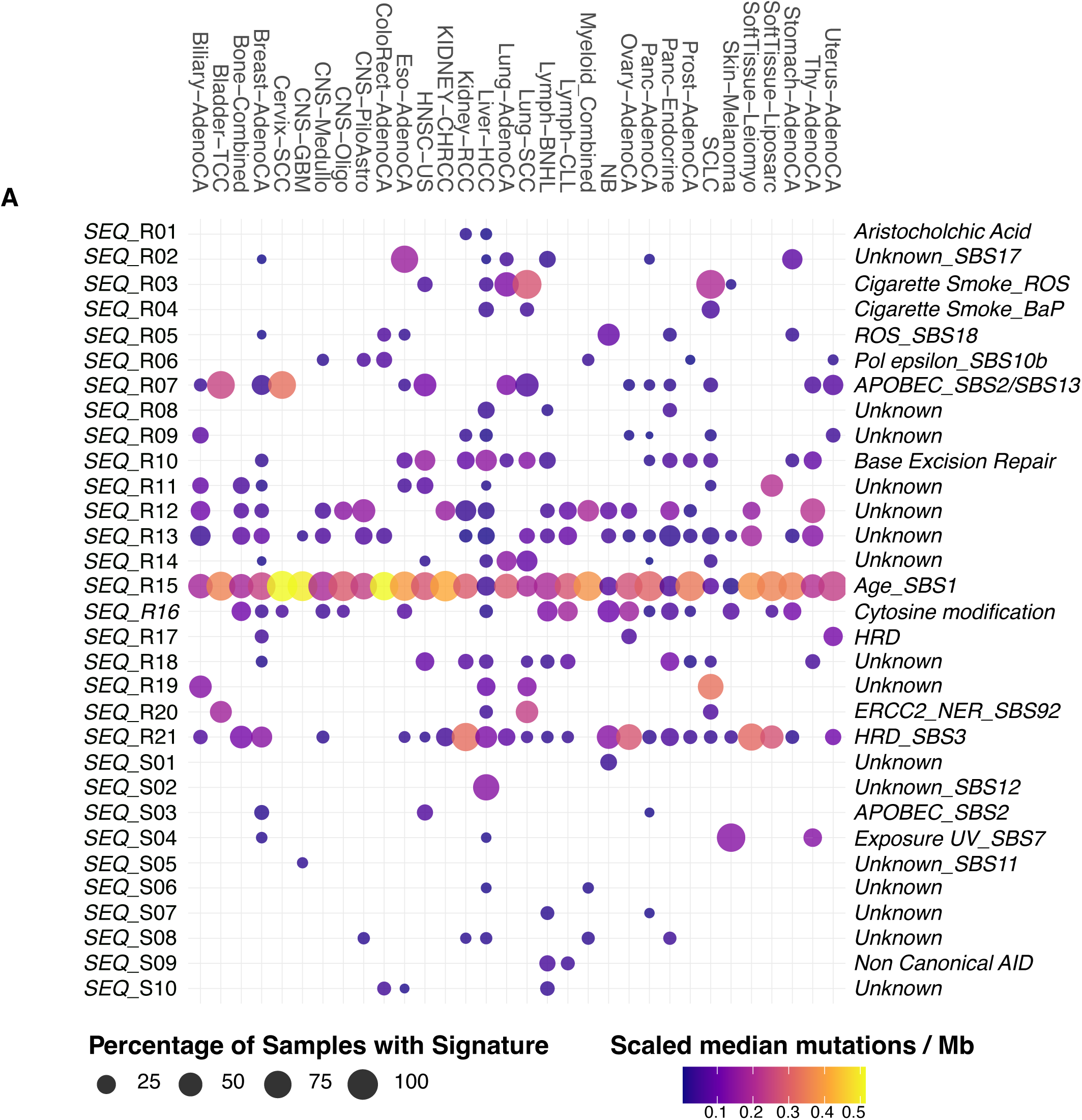
***SEQ* mutational signatures activities in the ICGC/PCAWG cohort.** A) Dot plot of *SEQ* mutational signatures activities in each tumour cohort and their putative etiology.

Association of signatures’ exposures with pathogenic genomic alterations in genes constituting the DNA repair pathway hint to mutation in the base excision repair machinery as the possible cause for *SEQ*_R10. The etiology of the remaining 13 signatures identified in this study is still unexplained (Suppl. Fig. 12).

## Discussion

Characterising the spectra of mutational processes has proven valuable for cancer prevention and clinical stratifications. Current analytical frameworks, which rely on matrix factorisation of mutation counts, implicitly assumes the availability of a given mutable sequence context uniform along the genome. However, the nucleotide composition of the human genome is far from homogeneous, as it is shaped by multiple evolutionary and molecular processes^46^. Furthermore, local mutation rates are dependent on chromatin organisation, replication timing, and transcription. Of note, DNA sequence and (epi)genomic states are interconnected and this influences mutational signature inference^13–15,20,46^. Thus, correcting mutation counts for the local availability of the sequence context in which they are observed to harmonise mutational opportunities across the genome, implicitly also minimises the influence of higher order of genomic organization on mutational signature extraction.

The scope of the present manuscript is to present *SEQ* as an accessible and flexible methodology to extract and interpret intrinsic mutation spectra of mutational processes in cancer, which are disentangled from the physical and physiological properties of the genome. Through an in-depth analysis of a series of mutational processes, we showed the main advantages of *SEQ* over the conventional aggregation of counts prior matrix factorisation. Thus, we show that mutational signature extraction upon *SEQ*: i) results in to generalised mutation spectra, less influenced by regional heterogeneity of the genome organisation, ii) provides clear tumour subtype classification, iii) disentangles co-occurring signatures with similar mutation spectra, iv) improves the inference of mutational signature with a flat spectrum, and v) can reveal novel aspects of mutagenesis. Furthermore, we publish a tentative reference set of mutational signatures generated upon *SE*Q. We are aware of its limitations and show it with the only purpose to reflect the reproducibility of our method in relation to the reference panel of mutational signatures published in Cosmic signatures. The analysis of additional whole genome tumour cohorts will enhance the breadth of the mutational signatures we have so far observed, refine reference mutational signatures that have been erroneously clustered, understand the etiology of some of the mutational signatures we have identified.

Some of the limitations of mutational signature inference are not addressed by *SEQ*: our method is not insensitive to tumour cohort size or signatures’ exposures. This explains our choice to infer mutational signatures on a tumour cohort basis rather than using combined datasets of samples across several tumour cohorts with unequal representation. In fact, flat mutational signatures and the extent of their exposures still affect the inference process^36^. This is further complicated by the fact that mutational processes often co-occur. While *SEQ* shows a substantial benefit in separating similar mutational spectra, we believe that disentangling process-specific effects remains a challenge^36,37^. The source code to generate *SEQ*-weighted mutation catalogues can be easily adapted to different genomes, genomic windows (e.g. for the analysis of whole exomes) as well as the sequence context categories. The current Cosmic and Signal databases of reference mutational signatures are extremely well curated with proven record to explain tumour etiology, cancer prevention and clinical stratification^6,7^. We believe that applying our method in future implementations of mutational signature extraction inference can improve the generalization and interpretation of mutational processes shaping the human genome.

## Methods

### Datasets used in this study

The ICGC/PCAWG cohort of 30 primary tumours (n=2638, EGAS00001001692)^16^, a small cell lung Cancer (n=101, EGAD00001001273)^17^ and a neuroblastoma (n=57, EGAS00001001308)^18^ cohort of WGS samples were used to infer mutational signatures based on *SEQ* mutational catalogues and create a reference dataset (Fig. 7, Suppl. Fig. 10a, Suppl. Fig. 11, Suppl. Table 1). The reference dataset was assigned to the Hartwig cohort of metastatic tumour samples (n=6135, www.hartwigmedicalfoundation.nl/en)^47^ to estimate the effect of the *SEQ*-HRD signature’s activity in predicting Homologous Repair Deficient tumours (see Fig.5a, Suppl. Table 1, and methods related to HRD). Signatures associated to cisplatin exposure were inferred from raw whole genome sequencing data of two cell lines, Hep2G and MCF10A, treated with a prolonged cisplatin treatment (Fig. 2 and Suppl. Fig. 2)^19^. Fastq files were obtained from the Eurpean Nucleotide Archive under the accession number PRJEB21971. Association of mutational signatures’ activity with 5-methyl cytosine (5-meC) was possible using deep whole genome bisulfite sequencing’ processed data derived from three lung bronchial tissues obtained from healthy donors. The data were downloaded from Gene Expression Omnibus using the accession numbers GSM5652337, GSM5652337, GSM5652337)^33^.

HRD-related signatures were inferred from the triple-negative breast cancer cohort SCAN-B (ClinicalTrials.gov ID NCT02306096)^48^. Somatic mutation data of 237 samples were downloaded from https://data.mendeley.com/datasets/2mn4ctdpxp/1. Patient data including the clinical outcome was obtained from^48^. Details on how the analysis of these data was performed can be found in methods section related to HRD and Fig. 5b and Suppl. Fig. 6. All data relative to the tubo-ovarian carcinoma University of Washington cohort -somatic mutation data, HRDetect scores and clinical data-described in Fig. 5c and Suppl. Fig. 7 was obtained through personal communication with Elisabeth M Swisher^35^. Details on how the analysis of these data was performed can be found in methods section related to HRD. Somatic mutation data to infer mutational signatures in healthy naive and memory B-cells (Fig. 6 and Suppl. Fig. 8 and Suppl. Fig. 9 ) were obtained from https://github.com/machadoheather/lymphocyte_somatic_mutation^45^. Processed bisulfite sequencing data of naive and germinal centre B-cells from wild-type and *Aicda ^-/-^*mice have been downloaded from the NCBI’s Gene Expression Omnibus under the accession number GSE71702 (Fig. 6h, 6i)^40^. Whole-genome bisulfite sequencing data of the lymphoblastoid human GM12878 cell line were downloaded from Encode with the accession number ENCFF279HCL and were used to estimate association between mutational signatures’ activities and 5-meC (Fig. 6g, Suppl. Fig. 8b). Details of how these data were analysed can be found in the methods section related to mutational processes of the B-cells.

### Opportunity-Weighted Mutation Counts

Regional somatic mutation density varies by more than an order of magnitude along the genome and depends on chromatin organization, replication timing and repair efficiency. Because these features are largely determined by cellular identity, the regional mutation density profile of a tumor retains an imprint of its cell of origin.

We propose that the same phenomenon has an underappreciated consequence for mutational signature analysis. Mutational signatures are conventionally defined as probability distributions over the 96 trinucleotide substitution contexts and are implicitly treated as fixed properties of the underlying mutational process, independent of the cellular context in which that process operates. However, the number of mutations observed in a given context depends not only on the intrinsic preference of the process but also on the mutational opportunity, that is, the number of available target sites for that context. Opportunity is commonly treated as a constant genome-wide (or exome-wide) trinucleotide composition, but this assumption is incomplete, because trinucleotide composition is itself regionally heterogeneous. The trinucleotide representation amplifies these differences, since each context occupies a narrower and more compositionally specific slice of sequence space than single nucleotides do. Taken together, these observations imply that the effective mutational opportunity is cell-type-specific. Each cell type has a characteristic landscape of chromatin accessibility and repair efficiency and therefore effectively weights different genomic regions, and hence different trinucleotide compositions, differently. Consequently, an identical mutational process acting in two different cell types is expected to produce systematically different 96-channel mutation spectra. This difference arises not from any change in the chemistry of the mutagenic process, but from differences in which parts of the genome are effectively available for mutation to occur and persist.

To correct for this, we replace the conventional mutation counts with opportunity-weighted counts, in which each mutation is weighted by the inverse of the local frequency of its trinucleotide, and perform signature extraction on the weighted catalogue. The resulting signatures are the intrinsic per-site mutation rates of each process, independent of the cell type and of the sequence composition of the analyzed territory.

### *De novo* inference of mutational signatures

Signatures of mutational processes causing somatic single nucleotide substitutions were inferred independently for each tumour type in each cohort using the *SigProfilerSuite* (v. 1.2.1, ref)^49,50,51^. The mutation catalogues used as input were either generated with *SEQUENCE* or with CaMuS’s module *matgen*^52^ or else, when indicated, with *SigProfilerMatrixGenerator*. *SEQ*-based mutation catalogues were obtained from the aggregation of mutated triplets weights calculated over sequential 10 kb genomic windows (see method section related to *SEQUENCE* implementation). All the signatures described in this work are the result of the *SigProfilerExtractor* function run in Python using default parameters’ values and setting the minimum_signature = 1 and the maximum_signature =7. In most instances, we used the number of mutational signatures suggested by the algorithm. Cosine Similarity and Kulback-Leibler divergence measures of the distance between the reconstructed matrices and the original catalogue were obtained from the *SigProfilerSuite* (Suppl. Fig. 10b, 10c).

### Methods related to mutational signatures inference of cisplatin treatment

To evaluate the ability of *SEQ* to extract cisplatin-associated mutational signatures across different cellular backgrounds, we analysed whole genome sequencing data from two cell lines, HepG2 and MCF10A, following a prolonged cisplatin treatment (see methods section related to datasets used in this study). Raw sequencing data were processed using the *PeifLyne* somatic mutation caller (http://www.uni-koeln.de/med-fak/peiflyne/peiflyne.tgz)^53^. Somatic mutations were then used to generate mutation catalogues and infer mutational signatures as described above. *Counts*-based mutational catalogues were obtained from filtered somatic mutations using *SigProfilerMatrixGenerator*.

### Assembly of a reference set of mutational signatures

To generate a reference dataset of mutational signatures, we applied hierarchical clustering with method="average" to the Euclidean distance matrix of pairwise 1-cosine similarities calculated among signatures. As general rule, a reference signature was assigned to a cluster of signatures when tree height was less or equal than 0.2 (Suppl. Fig. 10a). However, we acknowledge some mutational catalogues are more challenging to disentangle in to distinct mutational signatures than others, where factors such as number of samples, tumour mutation burden and signature activities penetrance can play a role in the stable inference of signatures of mutational processes. Therefore, manual adjustments to the afore mentioned rule was applied when necessary. Reference signatures (*SEQ*_R01 to *SEQ*_R21) were calculated as the average pattern across the singular mutational signatures within each cluster (Suppl. Fig. 10a, Suppl. Table 2). Ten signatures were not assigned to any cluster and they were labeled *SEQ*_S01 to *SEQ*_S10. Signature activities shown in Fig. 7 were obtained upon refitting the reference set to the ICGC/PCAWG samples. This procedure was performed stepwise: 1) each tumour cohort was fitted with the whole reference set of 31 *SEQ*-derived signatures; 2) a reference subset was defined based on signature activities present in at least four patients or signatures that had been inferred from a given cohort; 3) the reference subset was then refitted to each tumour cohort and the resulting activities are shown in Fig. 7. Signature etiologies, unless otherwise stated, were inferred by pattern similarities with the reference signature set published in Cosmic (https://cancer.sanger.ac.uk/signatures/) and prevalence of signature activities in given cancers.

### Association of mutational signatures with DNA repair pathway alterations

To investigate whether the *SEQ* reference mutational signatures are associated with defects in DNA repair pathways, we assessed the PCAWG samples for genomic alterations in selected DNA repair-associated driver genes. Genes were assigned to their respective DNA repair pathways, and a sample was classified as pathway-altered if at least one gene within the corresponding pathway was affected by a point mutation, copy number, or structural variant. Point mutations annotated as "pathogenic" in Clinvar or as "Drivers" in^7^, were considered for this analysis. Copy number alterations were defined as copy number losses (CN <2) in tumour suppressor genes or copy number amplifications (CN >=5) in oncogenes. Only structural variants whose breakpoint overlapped with the genomic coordinates of *MLH1* or *DDB2* were selected, given their negative association with the expression of these genes^54^. To account for differences in overall mutational signature activity between samples, signature activities were normalized to relative activities by dividing the absolute activity of each signature by the sum of all signature activities within the same sample. For each DNA repair pathway, samples classified as altered were compared to samples without alterations using Welch’s two-sample t-test to assess differences in mutational signature activities. Cohen’s d was calculated to quantify the standardized effect size, and p-values were adjusted for multiple testing using the Benjamini-Hochberg false discovery rate (FDR) procedure.

### Methods related to mutational signatures inference of cigarette smoking

To identify mutational signatures induced by the exposure to cigarette smoke in the ICGC/PCAWG cohort, we tested the inferred mutational signatures for their cosine similarity with mutational patterns obtained by experimentally exposing pluripotent stem cell to polycyclic aromatic hydrocarbons (PAHs)^27^. Variant call files were obtained from http://doi.org/10.17632/m7r4msjb4c.2. Specifically, we used samples MSM0.2 (BPDE 0.125 uM), MSM0.26 (BAP 0.39 uM), MSM0.42 (DBA 75 uM), MSM0.14 (DBADE 0.0313 uM), MSM0.12 (DBP 0.039 uM), MSM0.96 (DBPDE 0.000625 uM), MSM0.54 (5-Methylcrysene 1.6 uM), MSM0.74 (DBAC 5 uM). Mutation catalogues based on aggregated counts of mutation or based on *SEQUENCE* weighting were generated as described above. To identify mutational signatures of PAHs exposure in our cancer cohorts, we compared the mutation profiles obtained from pluripotent stem cells with the mutational signatures obtained from the SCLC, Lung AdenoCa, Lung-SCC, Head-Neck (the HNSC-US cohort, only smokers) and the Liver-HCC cohort, whose etiology is associated with cigarette smoking. We did so by initially clustering mutational signatures and mutational profiles based on their cosine similarities (method= "average", hclust () in R, library *stats*) and then selecting the PAH clade after manual inspection (Suppl. Fig. 4).

To compare dendrogram stability of the mutational signatures induced by exposure to PAH in cancers, we performed multinomial resampling (n=50) of the mutation catalogues (both *SEQ*-weighted and aggregated counts) obtained from the afore mentioned cancer cohorts. Each of the simulated matrices were inputted in *SigProfilerExtractor* to obtain mutational signatures. For each simulated dataset, mutational signatures were matched to their cognate reference signatures by cosine similarity and the PAH-induced simulated signatures were selected. The 50 simulated dendrograms of PAH-induced mutational signatures were compared to the original dendrogram using the prop.clades() in R from the *ape* library, returning how many times a node in the reference dendrogram is found in the simulated dataset.

Association of mutations driven by either *SEQ*_SCLC_SBS96A or *SEQ*_SCLC_SBS96C to 5-meC sites was tested using three lung bronchial tissues obtained from healthy donors and sequenced with deep whole genome bisulfite sequencing (see Datasets section). First, C>T somatic mutations in their respective triplet context were selected for contributing to either *SEQ*_SCLC_SBS96A or *SEQ*_SCLC_SBS96C based on an assignment probability ≥ 0.5. These mutations were then merged to methylation fractions based on their position along the chromosomes. Next, methylation fractions were binned and for each patient the number of mutations in each bin was counted and normalized to the total number of methylated sites within that bin. Comparison of mean mutation numbers between bin [0-0.09] and bin [0.9-1] was evaluated using a t-test.

### Methods related to mutational signature inference of Homologous Recombination-based Repair Deficiency (HRD) paragraph

To distinguish HRD from HRP samples we used HRDetect (Fig.5a left panel, Fig. 5b and Fig.5c)^5^ or CHORD (Fig.5a right panel)^55^ estimations. Samples with an HRDetect score higher or equal than 0.75 were classified as HRD and those with an HRDetect score lower or equal to 0.25 were classified as HRP. Samples with intermediate HRDetect scores, in the range between 0.26 and 0.74, were excluded from the analysis. Samples’ HRD assignment with CHORD in Fig. 5b was performed based on the "hr_status" variable being either "HR_proficient" or "HR_deficient".

#### Logistic regression

In order to identify mutational signatures associated to HRD, we performed logistic regression to test the association of signatures’ activities to samples classified as being either HRD or HRP. Signatures’ activities were log-transformed and z-score normalized to reduce the effect of outliers during the estimation process. We first performed a univariate screen so that for every candidate signature x_j_, we fit

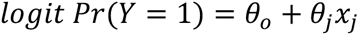

and retained the feature only if it reduced Akaike’s Information Criterion (AIC) relative to the intercept-only model, had a positive coefficient (θ^^^_j_ > 0). Starting from the intercept-only model, we then carried out forward stepwise selection on the set of retained signatures. Specifically, we iteratively fitted models adding a new signature at time and chose the signatures that yielded the lowest AIC (equivalently, the largest reduction in AIC from the current best model) subject to the positivity constraint θ^^^_j_ > 0. The procedure terminated when no remaining signatures decreased AIC. Final models were refitted with the selected signature set. All analyses were conducted in Python using *statsmodels* (GLM with binomial family, IRLS fitting), *pandas* for data handling, and *numpy* for numerical operations. For reproducibility, we logged, per tumour type, the null AIC, univariate screen results, the stepwise selection path (added features and AIC values), and the final model summary. The logistic regression has been used to aid the identification of the *SEQ*_R21 cluster (Suppl. Table 3), and the mutational signatures used in Fig. 5b and 5c.

#### Receiving Operator Characteristics (ROC) curve

ROC curves were generated to quantitatively compare the performance of *SEQ*_R21 and COSMIC SBS3 signatures’ activities in predicting HRD and HRP samples in the ICGC/PCAWG (Fig. 5a) and in the Hartwig cohort of metastatic tumours (Fig. 5b). For Fig. 5a we used HRD prediction from ^5^. Whereas for Fig.5b we used HRD prediction from ^55^. To obtain signature activities from mutation catalogues of aggregated counts we used *SigProfilerAssignment* and COSMIC_v3.6_SBS_GRCh37 as reference set. For *SEQ*-weighted mutation catalogues, *SigProfilerAssignment* was used together with our reference set (Suppl. Table 2). Signature assignment was performed stepwise similarly to the procedure described in the *Assembly of a reference set of mutational signatures* paragraph. Briefly, each tumour cohort was fitted with the whole reference set (i.e., COSMIC_v3.6_SBS_GRCh37 or Suppl. Table 2, depending on the experiment); next a reference subset was defined based on signature activities present in at least four samples; finally, the reference subset was refitted to each tumour cohort. ROC curves were generated using the *roc* function from the R library *pROC*. To test for significant differences in the Area Under the Curve (AUC) we used the *roc.test* function, method="delong".

#### Kaplan Meier (KM) analysis

To evaluate whether HRD-associated mutational signatures capture clinically meaningful differences, we performed Kaplan-Meier survival analyses in two prospective cohorts: the SCAN-B triple negative breast cancer^48^ and the University of Washington tubo-ovarian carcinoma cohort^35^. In both cohorts, mutational signatures were inferred using *SigProfilerExtractor*, with mutation catalogues derived from either aggregated counts or *SEQ*-based correction. Logistic regression (see above) was then applied to identify signatures associated with HRD using the available HRD scores (for details see Dataset paragraph). The activities of the identified signatures were log1p-transformed and z-score-standardized before patient stratification. Samples with signature activity ≥ their median value were classified as “HRD high”, and those < their median value as “HRD low”. When two signatures were considered jointly, samples were classified as “HRD high” if both activities were ≥ their respective medians and as “HRD low” if both were < their respective medians. Samples with discordant classification were excluded from that comparison. Mutational signature inference was performed from somatic mutation data of 237 samples of the triple negative breast cancer cohort (see paragraph *Dataset* for details). The HRD scores and the clinical variables for 165 samples in this cohort were obtained from the supplementary material of ^48^. Endpoints comprised invasive disease-free survival (IDFS), distant relapse-free interval (DRFI), and overall survival (OS). Event indicators were obtained from Clinical.IDFSbin, Clinical.DRFIbin, and OSbin (1 = event; 0 = censored), with corresponding times taken from Clinical.IDFS, Clinical.DRFI, and OS, measured in years. Samples were included in each endpoint analysis only when the corresponding analysis flag equaled 1 (Include IDFS, Include ACT DRFI Analysis, or Include ACT OS Analysis).

Mutational signature inference of the tubo-ovarian carcinoma cohort was performed on somatic mutation data of 185 samples (see paragraph *Dataset* for details). Recurrence/progression events were derived from RecurrenceOrProgression (Yes = 1; No = 0), with corresponding times taken from ProgressionFreeSurvival (PFS). Disease-specific survival events were derived from DiseaseStatus, with DOD coded as an event and AWD, NED, and D-other coded as censored, using the corresponding OverallSurvival times. Each endpoint analysis included only samples with an available survival time and a classifiable event status.

For both cohorts, survival distributions were estimated using the Kaplan-Meier estimator with right-censoring, and differences between strata were assessed using the two-sided log-rank test. Analyses were performed in Python using *lifelines* for survival estimation and log-rank testing, *pandas* for data handling and *matplotlib* for visualization.

### Methods related to signatures inference of the mutational processes operating in the B-cells

#### Inference of mutational signature in DLBCL super enhancers (SE)

In order to investigate AID-driven mutagenesis in diffuse large B-cell lymphoma, we subset the somatic variant call files of the DLBCL PCAWG cohort (histology tier4 Diffuse large B-cell lymphoma) based on the super enhancer coordinates retrieved from the supplementary materials in ^44^ (Figure 6a, 6b). For aggregated counts, SBS1536 mutation catalogues were generated using *SigProfilerMatrixGenerator*. To take in to account sequence bias due to selection of the super enhancer regions, relative context frequencies were calculated separately for the SE regions and the autosomal reference genome as

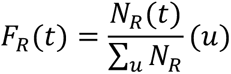

where N denotes the number of occurrences of pentanucleotide context (t) in sequence set (R), with (R) representing either the SE regions or the autosomal reference genome. Each mutation count was then corrected as

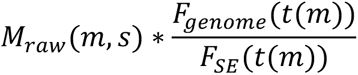

where (m) denotes the SBS1536 mutation channel, (s) denotes the sample, and (t(m)) is the reference pentanucleotide context corresponding to mutation channel (m). The same correction factor was applied to all substitution types sharing a reference context and to all samples. For contexts absent from the SE sequences, the correction factor was set to zero.

Annotation of the CLL samples as of whether originating from the germinal centre (i.e., MUT and UNMUT) was obtained from https://github.com/machadoheather/lymphocyte_somatic_mutation^45^.

To test the association of mutations driven by either *SEQ*_CLL_SBS96A or *SEQ*_CLL_SBS96B to methylated CpG sites we used methylation fractions derived from whole-genome bisulfite sequencing data of the GM12878 cell line (see Datasets section). This analysis was performed exclusively in CLL to ensure sufficient mutation counts for robust testing of each mutational signature. In fact, in healthy B-cells and DLBCL, several mutational signatures obtained after *SEQUENCE* correction are characterized by prominent C>T substitutions.

Positions of somatic mutation in LymphCLL were lifted to the reference genome GRCh38 using the liftOver pipeline by the USCS Genome Browser. Briefly, somatic mutations identified in CLL, along with their probabilities of assignment to either *SEQ*-CLL-SBS96A or *SEQ*-CLL-SBS96B, were mapped to the genomic positions in ENCFF279HCL. Methylation fractions were grouped into bins, and the number of methylated sites per bin was calculated. Somatic mutations were classified as associated with *SEQ*-CLL-SBS96A (probability ≥ 0.5) or *SEQ*-CLL-SBS96B. For each patient, the number of mutations per bin was counted and normalized by the total number of methylated sites within that bin. Comparison of mean number of mutations between bin [0-0.09] and bin [0.9-1] was evaluated using a t-test.

To obtain somatic mutations inside and outside the constant IG regions, we subsetted somatic variant call files based on the genomic coordinates of the constant region of the immunoglobulin genes (IGK-IGL-IGH). These files were then used to generate mutational catalogues transformed with *SEQ*. Finally, *SigProfilerAssignment* was used to assign mutational signatures to the mutational catalogues.

To test whether the AID-mediated deamination of the 5-meC is confined to WRCG DNA motifs, we obtained percentage of methylated and unmethylated sites from bisulfite sequencing data of B-cells derived from wild type and *Aicda -/-* mice. For each position, we extended the sequence context to +2 and -2 nucleotides surrounding the un(methylated) cytosines. We then classified the motifs as being WRCG or not-WRGC. In Fig. 6h, 6i, methylated sites refer to positions were 100% of the sequencing reads contain a cytosine, whereas unmethylated sites refer to positions were 100% of the sequencing reads contain a thymine.

## Acknowledgements

This work was supported by the German Research Foundation (DFG) through funding provided by a collaborative research centre grant on small cell lung cancer (CRC1399, project-ID 413326622) to MC, MP, JG, RKT, FB, MF and RHH. NA, MC, MP received additional fundings from the collaborative research centre grant on lymphoma B-cells (CRC1530, project-ID 455784452). JG, RKT, and MP received funding from the German Federal Ministry of Education and Research (BMBF, e:Med consortium InCa, grants 01ZX1901A and 01ZX2201A) and from "Netzwerke 2021", an initiative of the Ministry of Culture and Science of the German state of North Rhine-Westphalia, for the CANTAR project. Further support was provided by the DFG (project ID: 497777992 to JG, NA, MP, and AQ; project ID: GRK3110 to MP and JG), the German Cancer Aid (TACTIC, project ID 70115201, to RKT, and Mildred Scheel Professorship to MP), the Bruno-Helene-Joester Foundation (JG and MP), and the Jean Uhrmacher Foundation (JG). We furthermore thank the IT Center University of Cologne (ITCC) for providing computational resources on the DFG-funded high-performance computing clusters CHEOPS and RAMSES.

## Author Contribution

MC, NA, and MP conceived the project. MC, NA and FBierth wrote the manuscript. MC, NA and FBierth analysed the data. FBeleggia and RHH gave scientific input and contributed with data interpretation. JV and MS helped establishing the reference signature dataset. GP, CM worked on the experimental validation. RT and JG provided sequencing data of SCLC samples. MF provided sequencing data of neuroblastoma samples. MRR and ES provided genomic and clinical patients’ data of tubo-ovarian carcinoma samples. AP and MRS provided methylation data.

---

**Suppl. Fig. 1.**
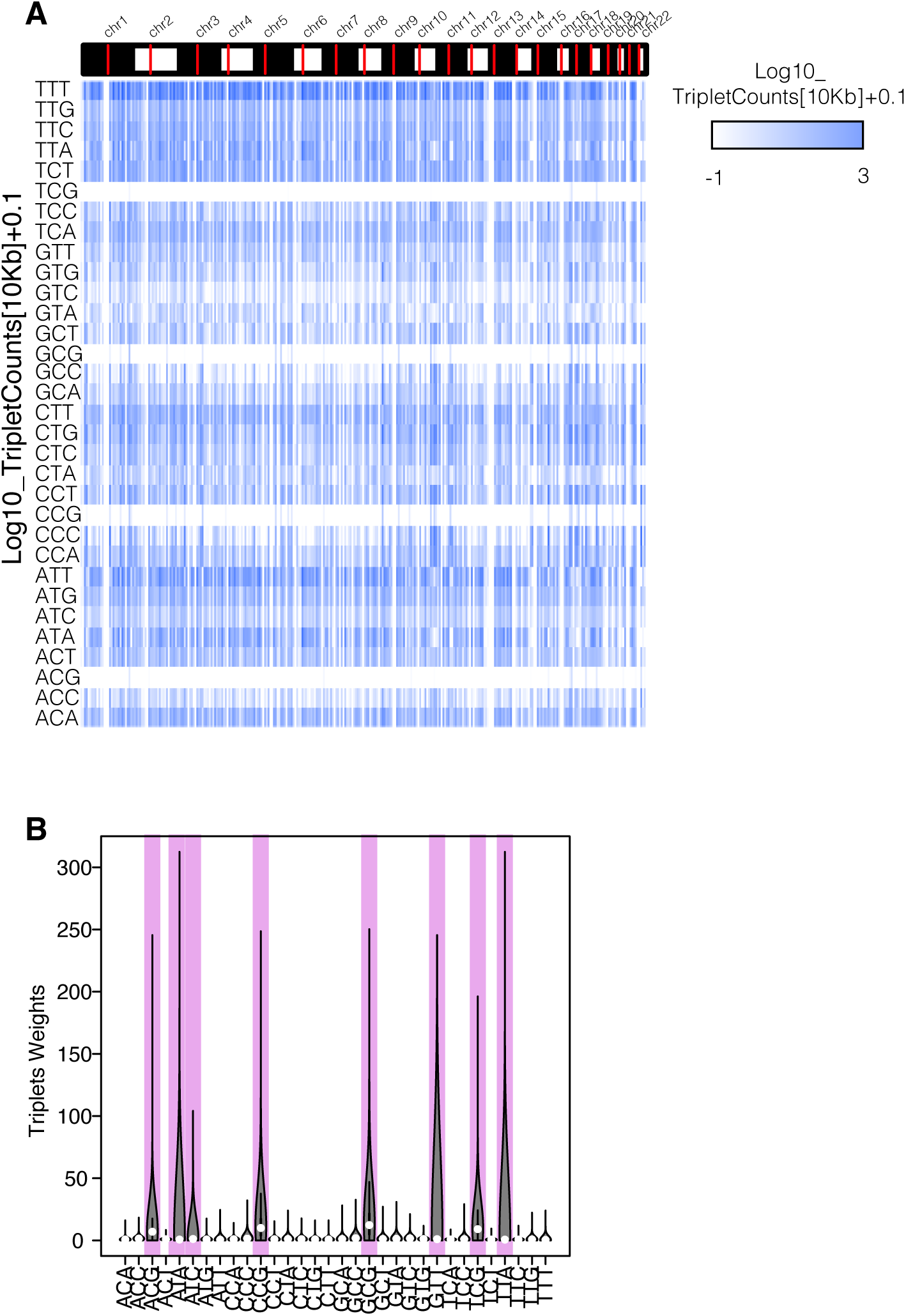
: Non uniform triplet distribution along the genome. A) Triplet counts within sequential-non-overlapping 10 kilobases windows. B) Boxplot of triplet weights estimated with *SEQ*. Pink boxes highlight triplets with highly heterogeneous weights.

**Suppl. Fig. 2.**
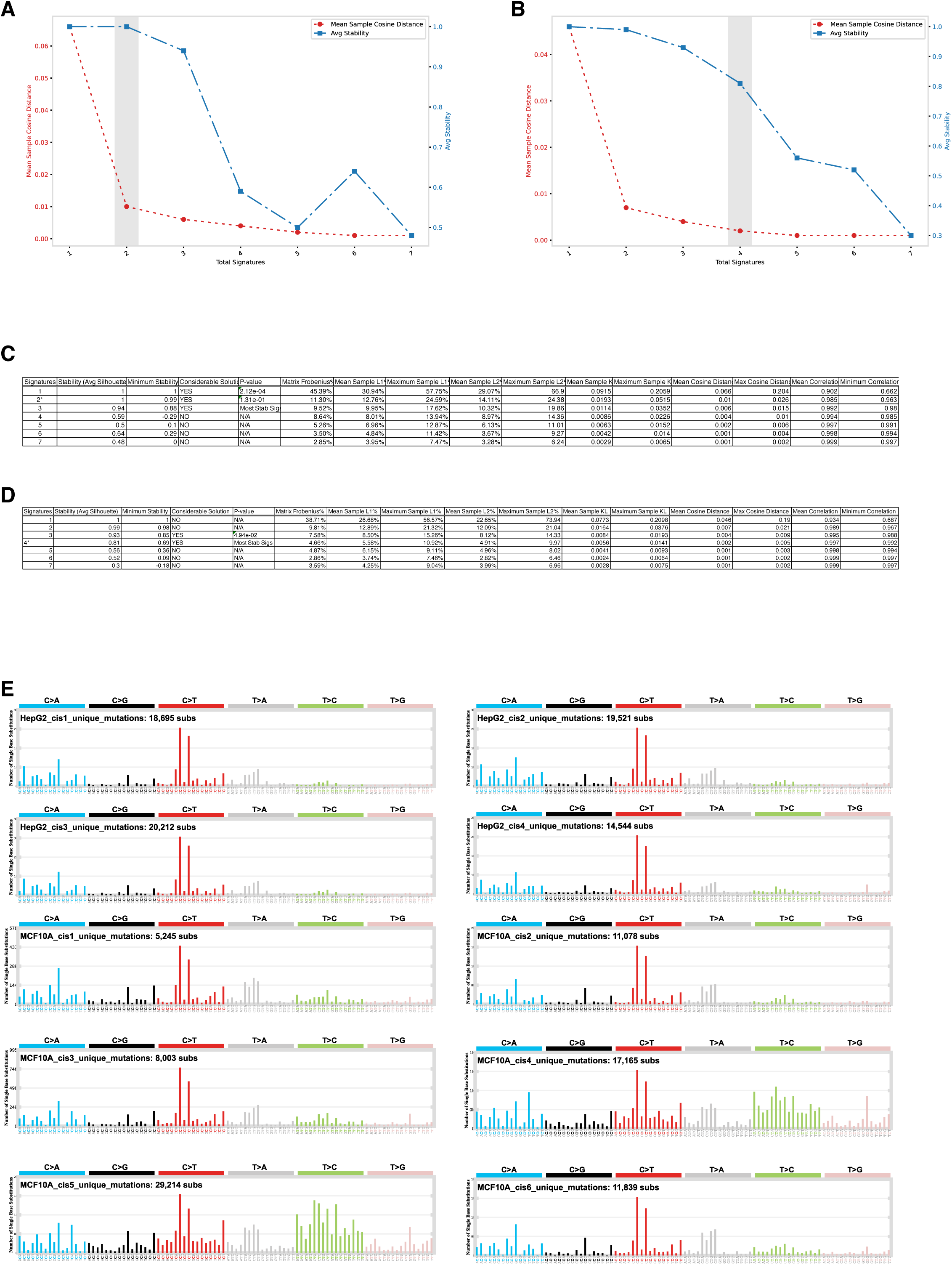
: Extraction of mutational signatures of cisplatin exposure using *SigProfilerExtractor*. Average signature stability (blue) and mean sample cosine distance (red) are plotted against the number of extracted signatures (1-7). The shaded grey bar indicates the optimal number of mutational signatures inferred using A) *SEQ*-weighted and B) raw *Counts* mutational catalogues as input. C) and D) Solution statistics for each extracted mutational signature upon either C) *SEQ* normalisation or D) raw *Counts*. E) Mutation profile of Hep2G and MCF10A cell line plotted against the 96 triplet combinations.

**Suppl. Fig. 3.**
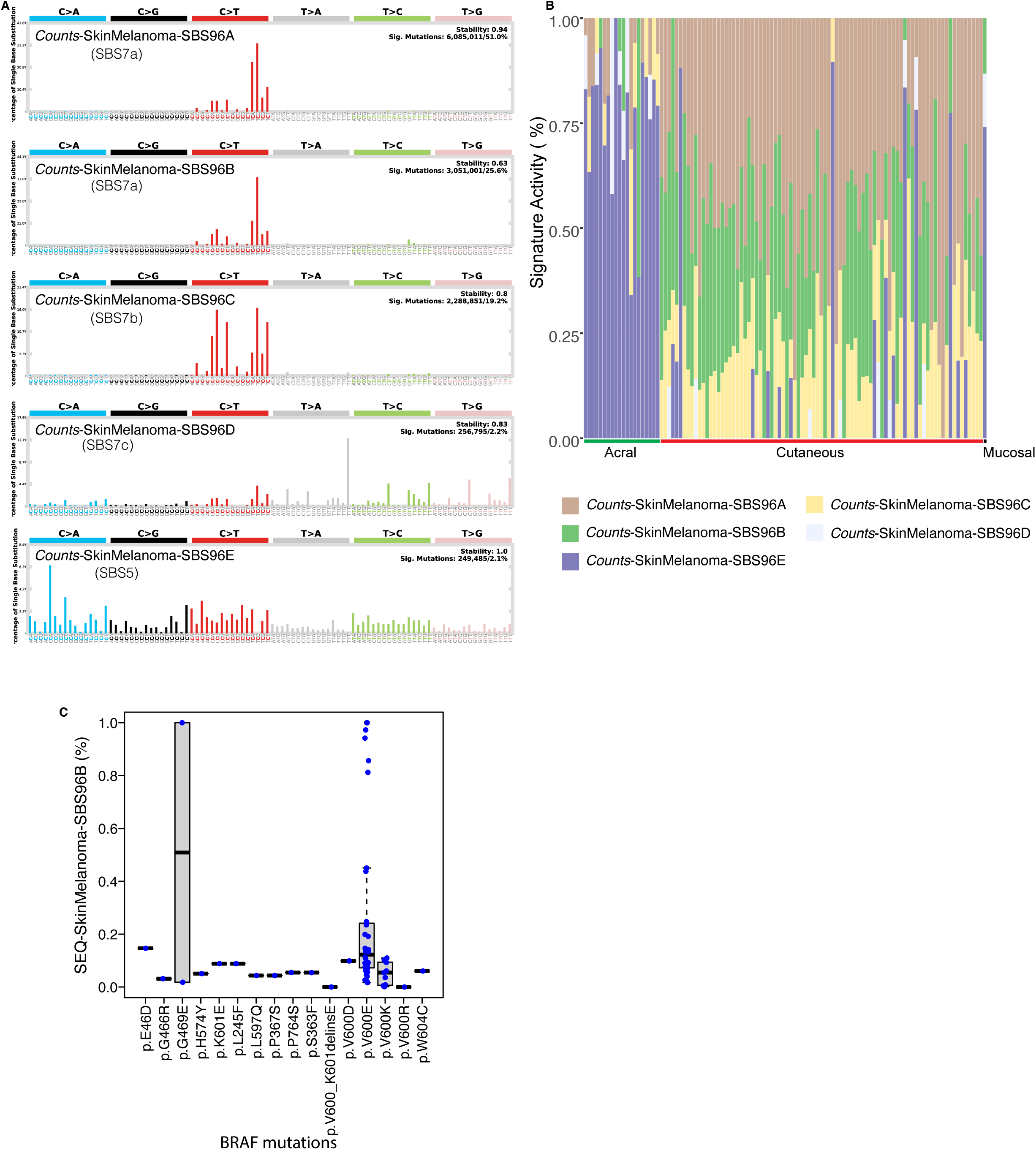
: *Counts*-derived mutational signature inference in the skin melanoma cohort. A) *Counts* mutational signatures profiles. Their similarity to the Cosmic reference panel set is also indicated. B) Stacked bar graph of relative mutational signature activity in the acral, cutaneous and mucosal subtypes. C) *SEQ*-SkinMelanoma-SBS96B activity in skin melanoma samples carrying *BRAF* mutant alleles.

**Suppl. Fig. 4.**
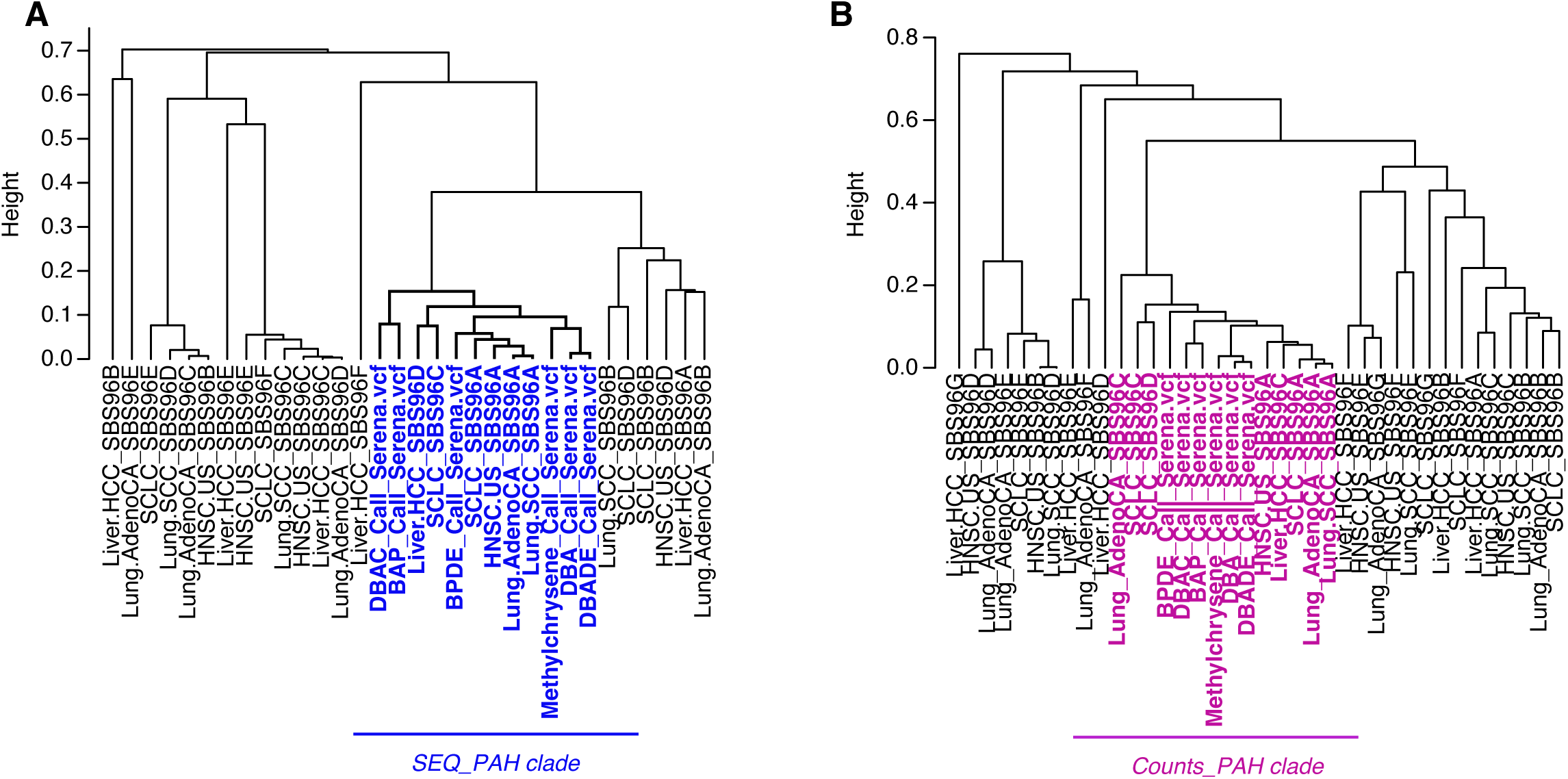
: Selection of mutational signatures derived from cigarette smoke exposure. A) Hierarchical clustering of 1-cosine pairwise similarities between PAH *SEQ*-weighted mutation catalogues and *SEQ*-weighted mutational signatures inferred in the Lung-SCC, Lung-AdenoCa, SCLC, Liver-HCC and HNSC-US cohorts. The estimated PAH clade is highlighted in blue for *SEQ* and in C) pink for *Counts*.

**Suppl. Fig. 5.**
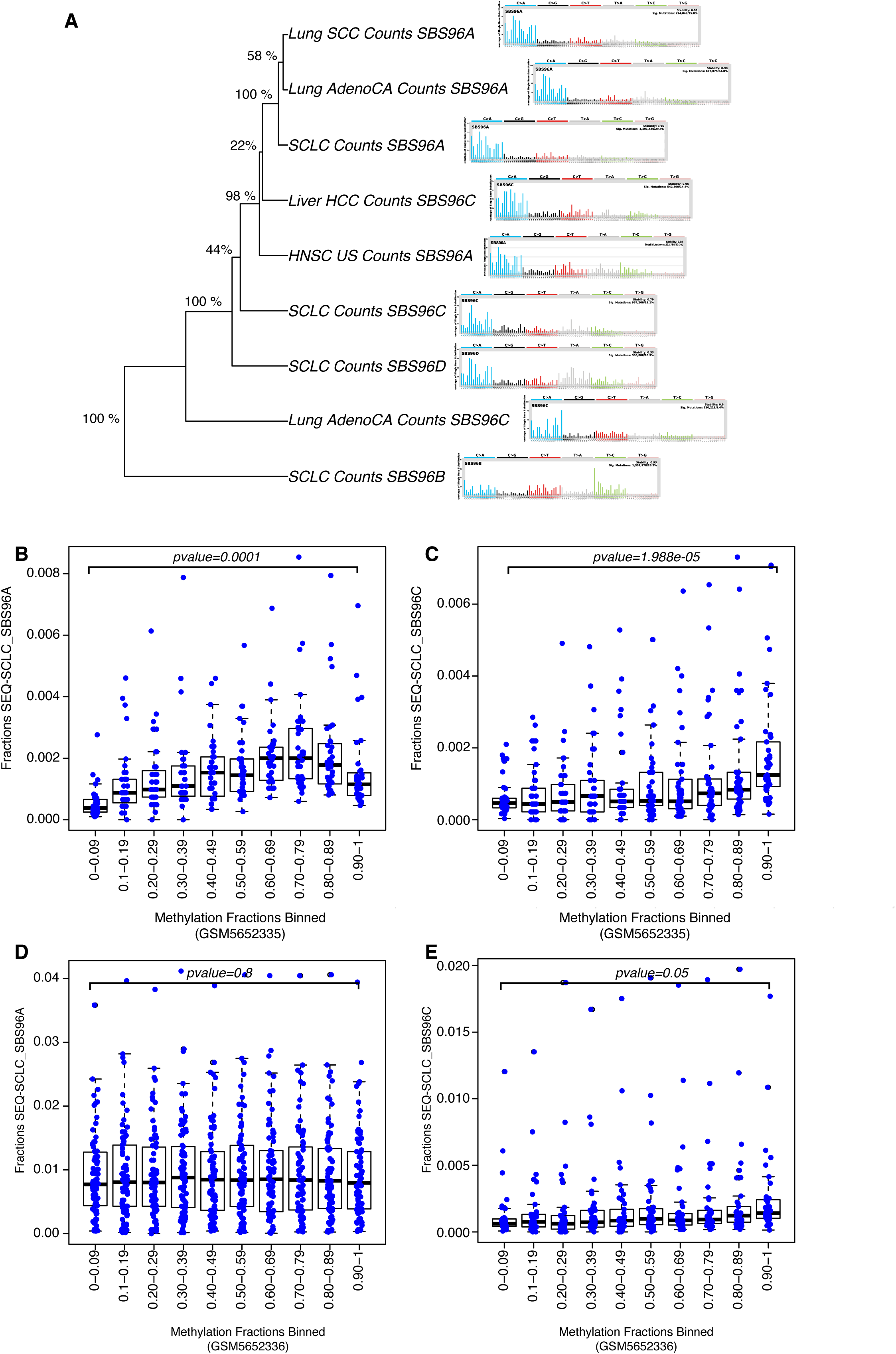
: *Counts*-derived mutational signatures of cigarette smoke exposure and association of *SEQ*-derived mutational signatures to methylated CpG sites. A) Hierarchical clustering of *Counts*-derived mutational signatures resulting from the exposure to cigarette smoke. The support for the branches is obtained from multinomial resampling of the original mutational catalogues and it is shown as percentage on the left of the tree. B) and D) Fractions of mutations assigned to *SEQ*-SCLC_SBS96A at methylated sites measured in a lung bronchial tissue cell lines from healthy donors (GSM5652335 and GSM5652336, respectively). C) and E) Fractions of mutations assigned to *SEQ*-SCLC_SBS96C at methylated sites measured in a lung bronchial tissue cell lines from healthy donors (GSM5652335 and GSM5652336, respectively). Methylation fractions are separated in to bins. P-values obtained from student’s t-test comparing fractions in the first (0-0.09) and the last bin (0.90-1) are indicated on the top of the boxplots.

**Suppl. Fig. 6.**
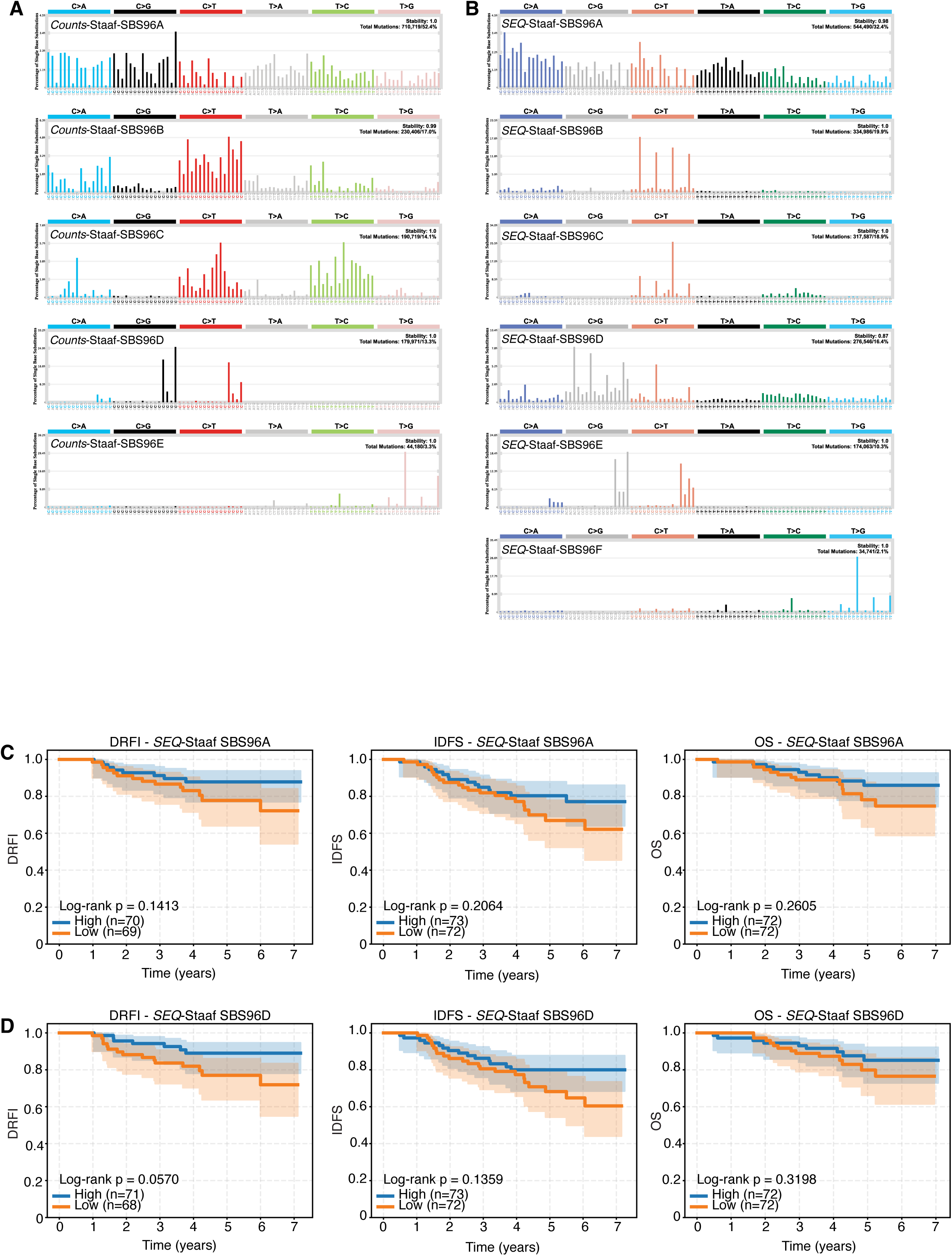
: Mutational signature inference in the triple negative breast cohort *Staaf et al*. Mutational signatures inferred with *Counts*-based (A) and *SEQ*-weighted mutational catalogues (B). C) Kaplan Meier curves and log rank test reflecting the ability of the *SEQ*_Staaf-SBS96A signature to stratify patients by DRFI, IDFS and OS. D) Kaplan Meier curves and log rank test reflecting the ability of the *SEQ*_Staaf-SBS96D signature to stratify patients by DRFI, IDFS and OS. Orange and dark blue indicate patients stratified according to the activity of the HRD signatures estimated with *SEQ*. High and Low indicate above/equal and below median signature activity, respectively. n= number of patients belonging to the above and below the median groups, respectively. DRFI: Disease Relapse Free Interval, IDFS: Invasive Disease-Free Survival, OS: Overall Survival.

**Suppl. Fig. 7.**
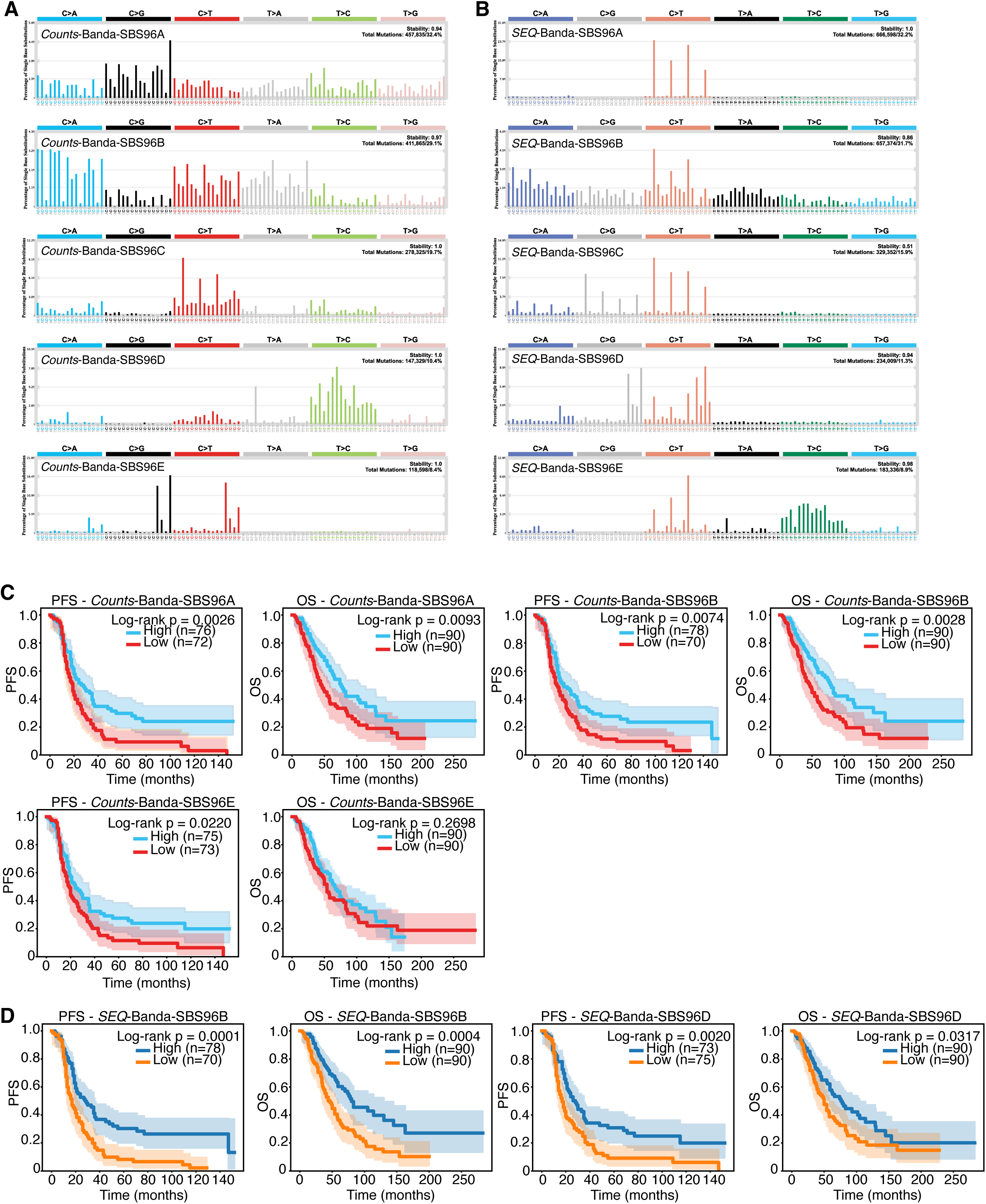
: Mutational signature inference in the tubo-ovarian carcinoma Washington cohort *Banda et al*. Mutational signatures inferred with *Counts*-based (A) and *SEQ*-weighted mutational catalogues (B). C) Kaplan Meier curves and log rank test reflecting the ability of the *Counts*_Banda-SBS96A, *Counts*_Banda-SBS96B and *Counts*_Banda-SBS96E signatures alone to stratify patients by PFS and OS. D) Kaplan Meier curves and log rank test reflecting the ability of the *SEQ*_Banda-SBS96B and *SEQ*_Banda-SBS96D signatures alone to stratify patients by PFS and OS. Orange and dark blue indicate patients stratified according to the activity of the HRD signatures estimated with *SEQ*. High and Low indicate above/equal and below median signature activity, respectively. n= number of patients belonging to the above and below the median groups, respectively. OS: Overall Survival, PFS: Progression Free Survival. Red and light blue indicate patients stratified according to the activity of the HRD signatures estimated with counts. Orange and dark blue indicate patients stratified according to the activity of the HRD signatures estimated with *SEQ*. High and Low indicate above/equal and below median signature activity, respectively.

**Suppl. Fig. 8.**
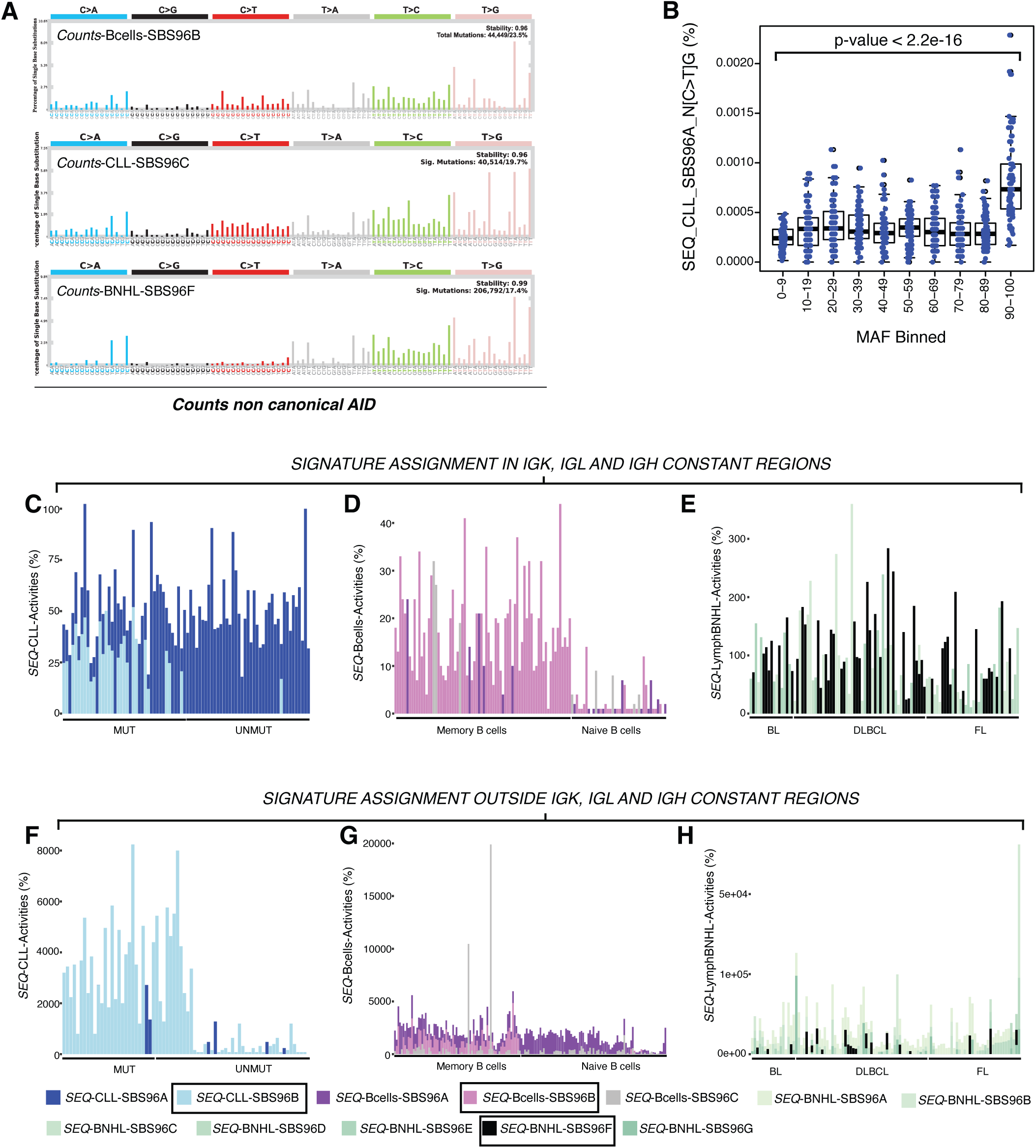
: Mutational signatures of non-canonical AID in B-cells. A) *Counts*-based mutational signatures of non-canonical AID independently inferred in a cohort of healthy memory and naive B-cells, Lymph-CLL and Lymph-BNHL tumours. B) Association of the N[C>T]G mutations assigned to *SEQ*-CLL_SBS96A to methylation allelic frequencies in the lymphoblastoid cell line GM12878. P-value of a student’s t-test between bins (0-9) and (90-100) is indicated on top. Assignment of mutational signature activities within the *IGK*, *IGL* and *IGH* constant region in Lymph-CLL C), healthy B-cells D) and Lymph-BNHL E). Assignment of mutational signature activities outside the *IGK*, *IGL* and *IGH* constant and switch region in Lymph-CLL F) healthy B-cells G) and Lymph-BNHL H). MUT indicates Lymph-CLL tumours originating from the germinal centre. UNMUT indicates Lymph-CLL tumours originated before entering the germinal centre. FL is Follicular Lymphoma, BL is Burkitt Lymphoma and DLBCL is Diffuse Large B-cell Lymphoma.

**Suppl. Fig. 9:**
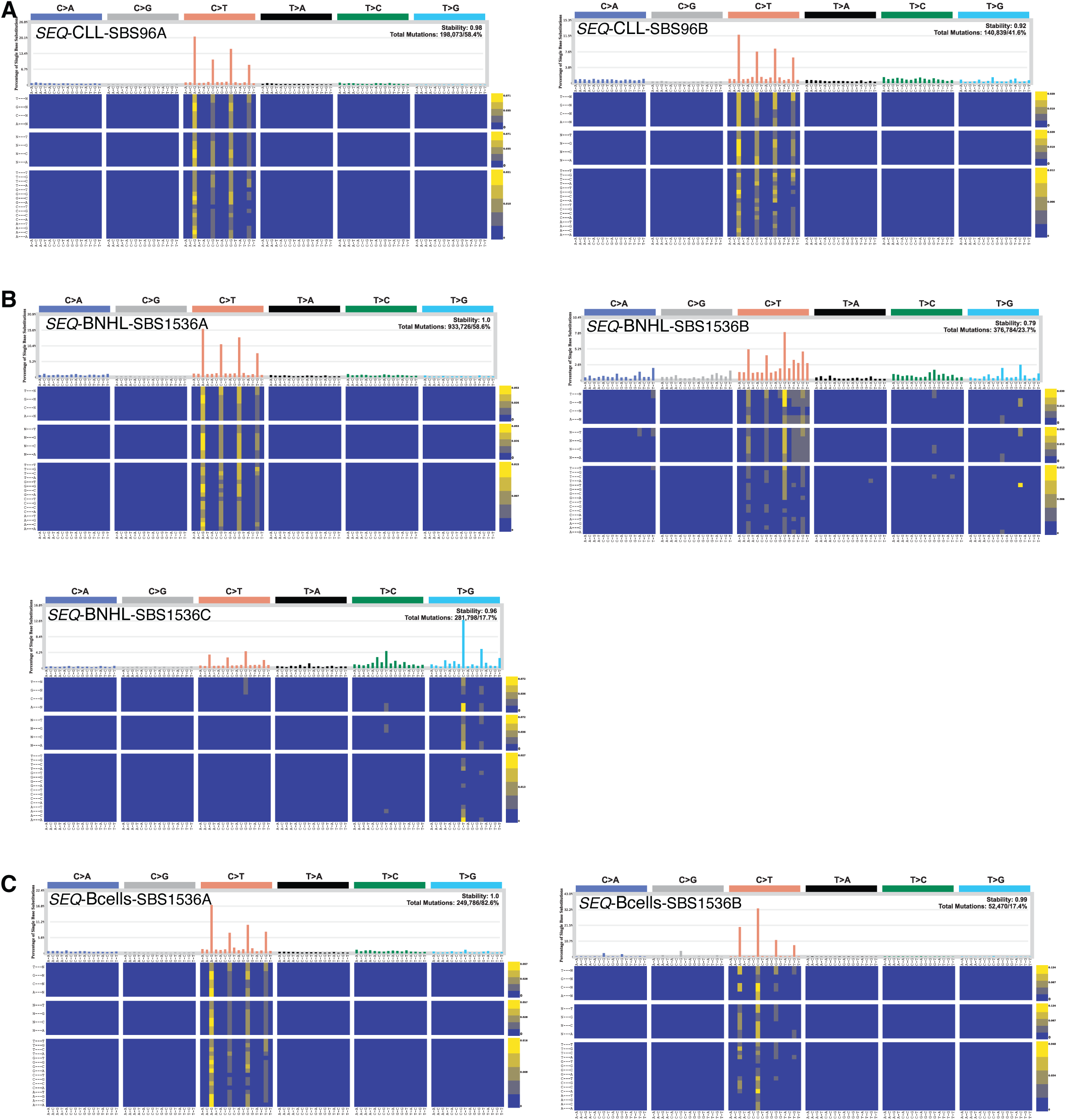
*SEQ*-based SBS1536 mutational signatures in lymphomas and healthy B-cells. Mutational signature extraction in Lymph-CLL A); Lymph-BNHL B); healthy B-cells C).

**Suppl. Fig. 10:**
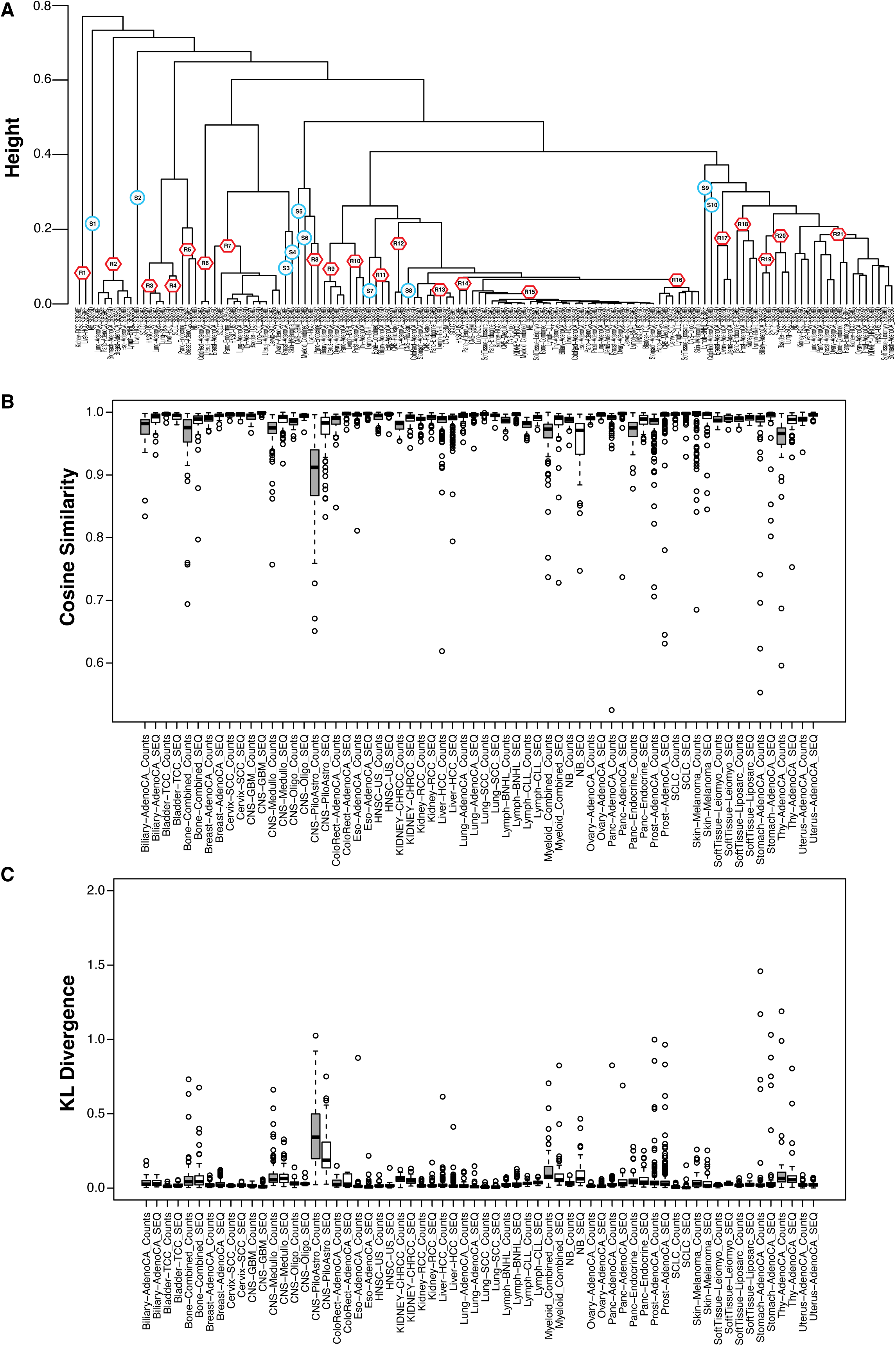
*SEQ* mutational signatures inference in the ICGC/PCAWG, SCLC and Neuroblastoma cohort. A) Hierarchical clustering of mutational signatures based on pairwise 1-cosine similarities between their mutation profile. Red hexagons indicate clusters of signatures exhibiting similar mutation profiles. Blue circles indicate signatures whose mutation profile is substantially different from those of other signatures. Cluster membership was determined upon manual curation using the dendrogram height <=0.2 as rule of thumb. B) Cosine Similarities and C) KL divergence between the reconstructed and the original mutational catalogues, derived using either with *SEQ* or with raw *Counts*.

**Suppl. Fig.11:**
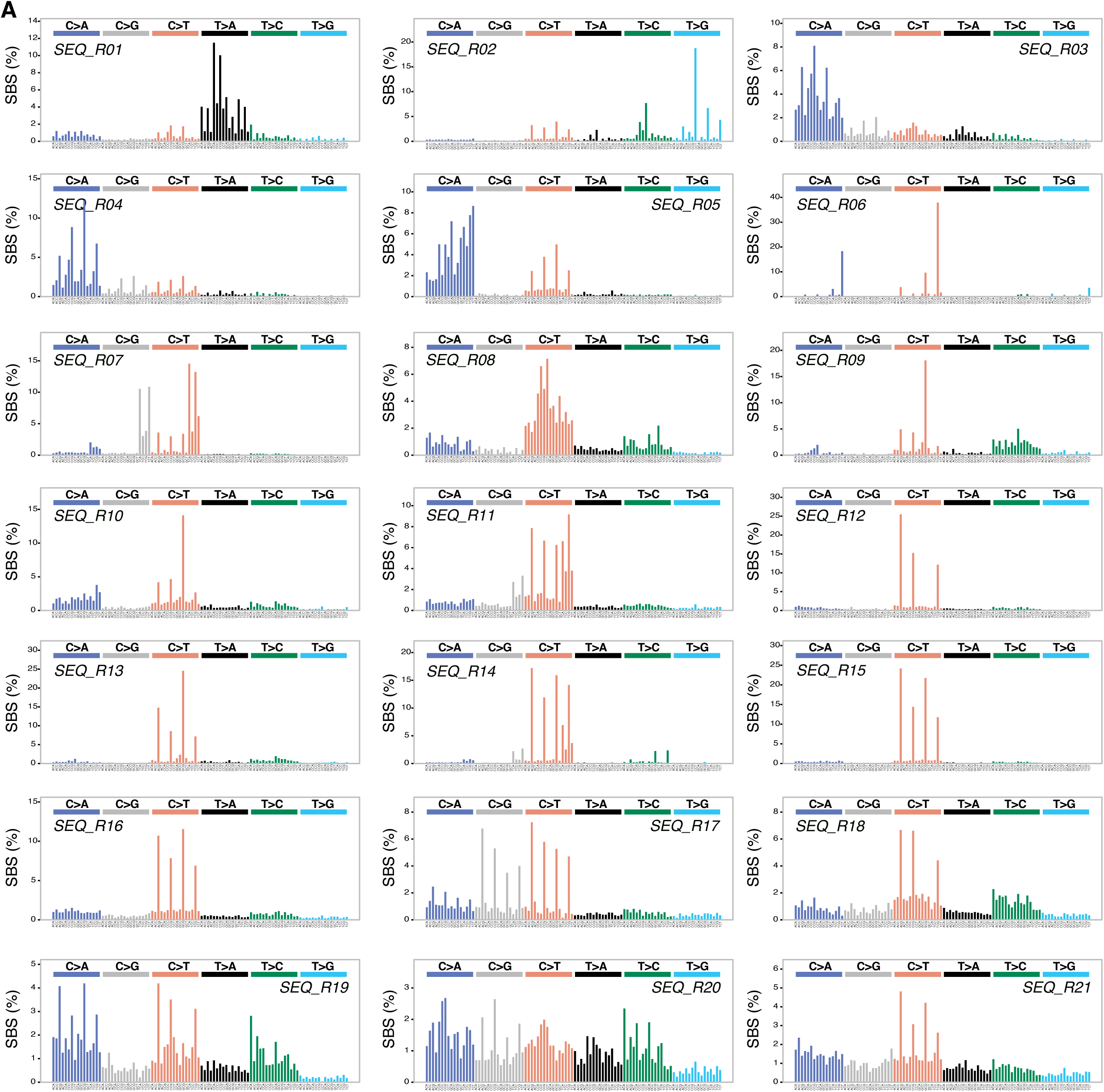
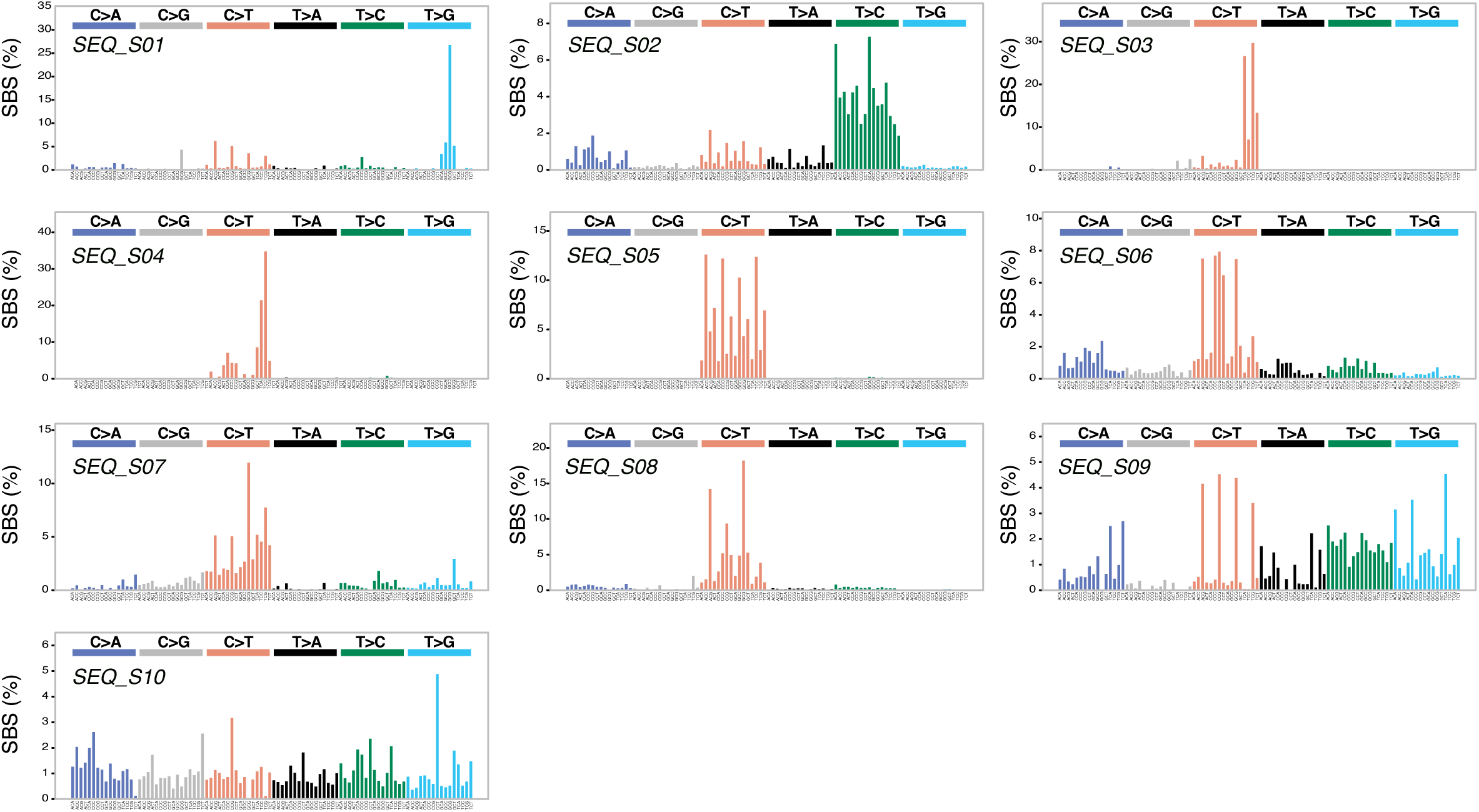
Mutation profiles of *SEQ*-derived reference mutational signatures.

**Suppl. Fig. 12.**
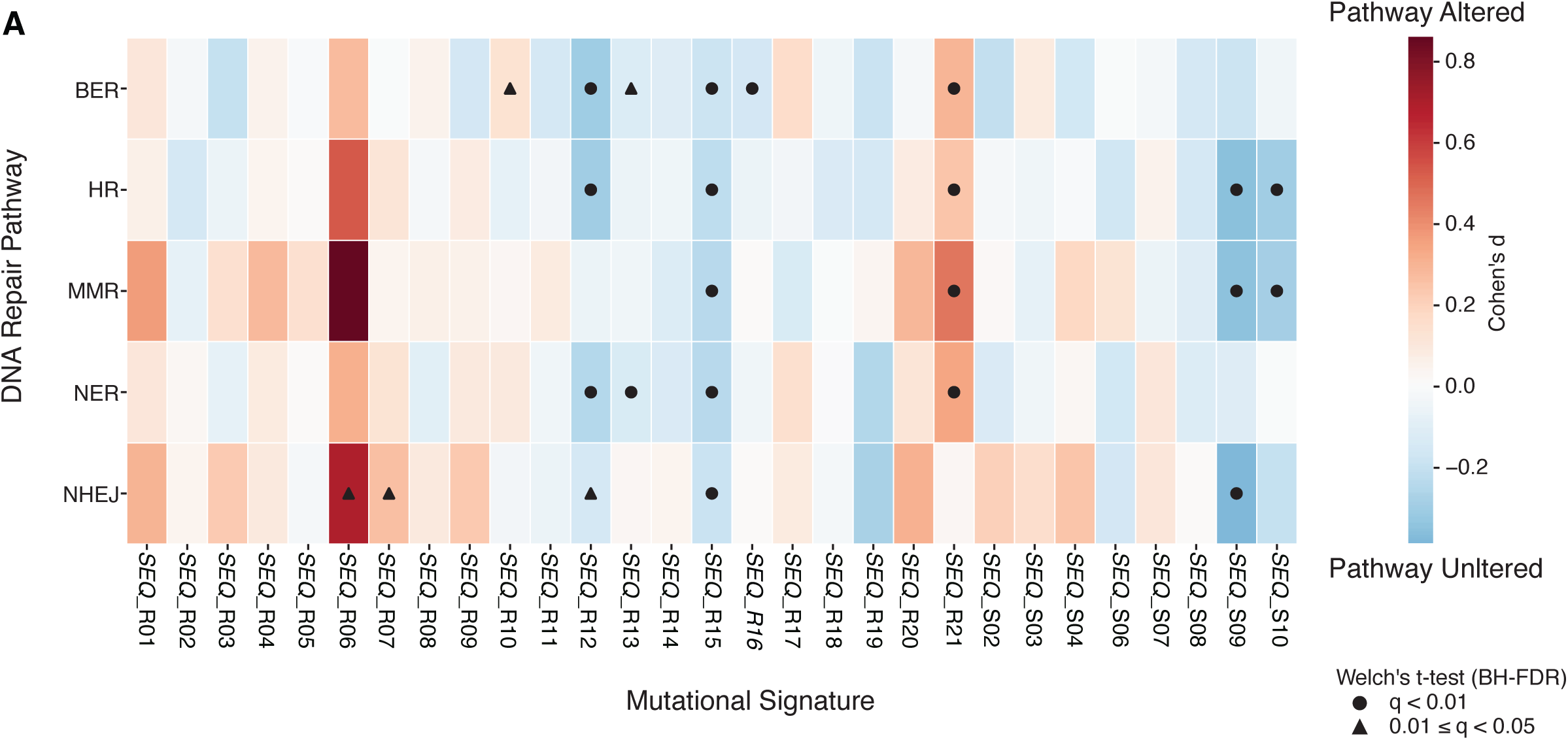
: Association between *SEQ-derived* mutational signatures activities and somatic alteration in the DNA repair machinery.

